# RhizoVision Crown: An Integrated Hardware and Software Platform for Root Crown Phenotyping

**DOI:** 10.1101/569707

**Authors:** Anand Seethepalli, Haichao Guo, Xiuwei Liu, Marcus Griffiths, Hussien Almtarfi, Zenglu Li, Shuyu Liu, Alina Zare, Felix B. Fritschi, Elison B. Blancaflor, Xue-Feng Ma, Larry M. York

**Affiliations:** Noble Research Institute, LLC, 2510 Sam Noble Parkway, Ardmore, OK 73401; Division of Plant Sciences, University of Missouri, Columbia, MO, 65201; Crop and Soil Sciences, University of Georgia, Athens, GA, 30602; Texas A&M AgriLife Research, Texas A&M University System, Amarillo, TX, 79106; Department of Electrical and Computer Engineering, University of Florida, Gainesville, FL, 32601

## Abstract

Root crown phenotyping measures the top portion of crop root systems and can be used for marker-assisted breeding, genetic mapping, and understanding how roots influence soil resource acquisition. Several imaging protocols and image analysis programs exist, but they are not optimized for high-throughput, repeatable, and robust root crown phenotyping. The RhizoVision Crown platform integrates an imaging unit, image capture software, and image analysis software that are optimized for reliable extraction of measurements from large numbers of root crowns. The hardware platform utilizes a back light and a monochrome machine vision camera to capture root crown silhouettes. RhizoVision Imager and RhizoVision Analyzer are free, open-source software that streamline image capture and image analysis with intuitive graphical user interfaces. RhizoVision Analyzer was physically validated using copper wire and features were extensively validated using 10,464 ground-truth simulated images of dicot and monocot root systems. This platform was then used to phenotype soybean and wheat root crowns. A total of 2,799 soybean (*Glycine max*) root crowns of 187 lines and 1,753 wheat (*Triticum aestivum*) root crowns of 186 lines were phenotyped. Principal component analysis indicated similar correlations among features in both species. The maximum heritability was 0.74 in soybean and 0.22 in wheat, indicating differences in species and populations need to be considered. The integrated RhizoVision Crown platform facilitates high-throughput phenotyping of crop root crowns, and sets a standard by which open plant phenotyping platforms can be benchmarked.

## INTRODUCTION

Roots serve as the interface between the plant and the complex soil environment with key functions of water and nutrient extraction from soils (Lynch, 1995; Meister et al., 2014). Root system architecture (RSA) refers to the shape and spatial arrangement of root systems within the soil, which plays an important role in plant fitness, crop performance, and agricultural productivity (Lynch, 1995; York et al., 2013; Rogers and Benfey, 2015). RSA is shaped by the interactions between genetic and environmental components, and it influences the total volume of soil that roots can explore (Rogers and Benfey, 2015). Many root phenes (or elemental units of phenotype Lynch, 2011; Pieruschka and Poorter, 2012; York et al., 2013) shape the final RSA, including the number, length, growth angle, elongation rate, diameter, and branching of axial and lateral roots (Bishopp and Lynch, 2015). Understanding the contribution of RSA phenes to crop performance is of key importance in food security and for breeding of more productive and resilient varieties in a changing environment.

Because roots are hidden underground and require considerable effort to characterize, research on roots lags behind that on aboveground parts of the plant (Eshel and Beeckman, 2013), and the genetic and functional basis of RSA remains obscured (Topp et al., 2016). Phenotyping is a major bottleneck in research and a lack of efficient methods for collecting root phenotypic data is limiting progress in using RSA for genetic studies and breeding for root ideotypes (Das et al., 2015; Kuijken et al., 2015). In recent years there has been a shift to image-based phenotyping for enabling relatively high-throughput and accurate measurements of roots. Many of the platforms use 2D imaging with cameras, and involve the use of seedlings on agar plates, germination paper or fabric cloth in bins (Kuijken et al., 2015). Despite the usefulness of controlling environmental parameters for characterization of root phenotypes, studies of roots of field-grown plants better represent the agricultural systems in which they ultimately grow.

Weaver and colleagues (Weaver, 1925; Weaver and Bruner, 1926) pioneered methods for excavating, drawing and photographing root systems that have been widely used for more than half a century (Böhm, 2012). These classical methods were since modified (Stoeckeler and Kluender, 1938) with the use of water to remove soil particles from the root systems on a large scale, and using high pressure air to penetrate soil pores while leaving roots intact (Kosola et al., 2007). Hydropneumatic root elutriation is a different method developed by Smucker et al. (1982) to provide a rapid and reproducible approach for separating roots from soil of field-collected soil core samples with minimal damage. Traditional excavation methods are most suited for trees and shrubs as the root system of wooden species are generally stronger and more resistant to breaking than the finer roots of grasses or annual crops (Böhm, 2012). Other field root phenotyping methods include minirhizotrons and soil coring, which both require a large amount of physical labor and set-up time (Johnson et al., 2001; Böhm, 2012; Wasson et al., 2016). More recently non-destructive root phenotyping methods such as ground penetrating radar and electrical resistance tomography have shown promise, however both techniques only provide indirect assessments of root length and do not provide RSA (Garré et al., 2013; Liu et al., 2018).

Over the past 10 years, root crown phenotyping (York, 2018) has emerged as one of the more common field-based root phenotyping methods, and is characterized by excavation of the top portion of the root system, removal of soil, and measurements, by a variety of means. The definition of root crown as the top portion of the root system in this research is extended from the earlier use of this terminology which refers to the site where the root system transitions to the shoot (Beentje, 2010). Root crown phenes, such as nodal root number (York et al., 2013; Gao and Lynch, 2016; Slack et al., 2018) and growth angle (Wasson et al., 2012; Trachsel et al., 2013; York et al., 2015; Slack et al., 2018), have been widely reported to correlate with crop above-ground biomass or grain yield. The work of Grift et al. (2011) may be the earliest published example of root crown phenotyping in a high-throughput capacity. Root crown phenotyping was widely popularized as “shovelomics” in the work of Trachsel et al. (2011) using visual scoring. While the term “shovelomics” is popular, the extent of its definition is not clear and debate exists whether it only refers to methods based on root crown washing and visual scoring in maize (*Zea mays* L.) or to other protocols. Therefore, “root crown phenotyping” is proposed as less ambiguous and more broadly applicable, as defined above. Root crown phenotyping has been used to enhance the understanding of soil resource acquisition by roots of soybean (*Glycine max* L.), common bean (*Phaseolous vulgaris* L.), cowpea (*Vigna unguiculata* L.), wheat (*Triticum aestivum* L.), and maize (Trachsel et al., 2010; Colombi et al., 2015; York et al., 2015; York and Lynch, 2015; Burridge et al., 2016; Maccaferri et al., 2016; York et al., 2018).

In order to standardize measurements and increase throughput, image-based phenotyping of crop root crowns has become the standard procedure. The unique steps of image-based phenotyping are acquiring and analyzing the image, which are of equal importance with regards to creating a reproducible and reliable protocol. Potentially, the first example of image-based root crown phenotyping used a custom imaging booth with vision cameras controlled by MATLAB and image analysis in MATLAB that provided two measures, fractal dimension and top root angle (Grift et al., 2011). The Digital Imaging of Root Traits (DIRT) platform attempted to relax imaging requirements by allowing use of any consumer camera with roots generally placed on a dark background in uncontrolled lighting conditions and currently focuses on a free cloud-based image analysis pipeline, though a Linux installation is possible (Bucksch et al., 2014; Das et al., 2015). The Root Estimator for Shovelomics Traits (REST) platform included an imaging ‘tent,’ DSLR consumer camera controlled using the manufacturer’s software, and a MATLAB executable for image analysis (Colombi et al., 2015). The Multi-Perspective Imaging Platform (M-PIP) includes five point-and-shoot cameras along a 90° arc in an imaging box, command line camera control software for Linux, and MATLAB scripts for image analysis (Seethepalli et al., 2018). The cloud-based platform of DIRT requires uploading potentially thousands of root images, which is time consuming, and then downloading the data, and the less-controlled imaging protocol leads to segmentation failures. The REST platform provides controlled imaging conditions, though not with optimal ergonomics, and the MATLAB implementation doesn’t include root length. M-PIP requires knowledge of Linux and requires difficult segmentation of roots form the background using color information. These platforms have advanced the field of root crown phenotyping, but advances can still be made to increase access to these technologies and to optimize imaging, image analysis, and data processing.

The aim of this study was to develop a phenotyping platform for both high-throughput image acquisition and image analysis of root crowns from the field. The imaging hardware is ergonomic and portable for the user, reproducible in any lab, and affordable. The imaging software is optimized for rapid plant phenotyping and usability. The image analysis software is designed to be extremely fast, reliable, fully automated. Several validation tests were performed with the software which have achieved a 100% success rate when used with images from the hardware platform. Together, these developments represent an elegant solution for root crown phenotyping.

## MATERIALS AND METHODS

### Experimental Design

In order to achieve the goals to design and validate a new phenotyping platform for crop root crowns, several related tasks and experiments were conducted (Fig. 1). The backlit RhizoVision Crown hardware platform was designed and tested. The RhizoVision Imager program was developed to interface with the camera of the hardware platform and RhizoVision Analyzer was developed for processing images of root crowns to generate tabular and numeric data for further analysis. Validation of the platform’s ability to generated accurate physical measurements was done using scans of copper wires of known diameter. In order to validate root measurements such as length, area, and number of root tips, simulated datasets of monocot and dicots were used. In order to test the hardware platform, imaging software, and analysis software, root crown phenotyping was conducted for a soybean population in Missouri and a wheat population in Oklahoma. These experiments are discussed in greater detail below.

**Figure 1.**
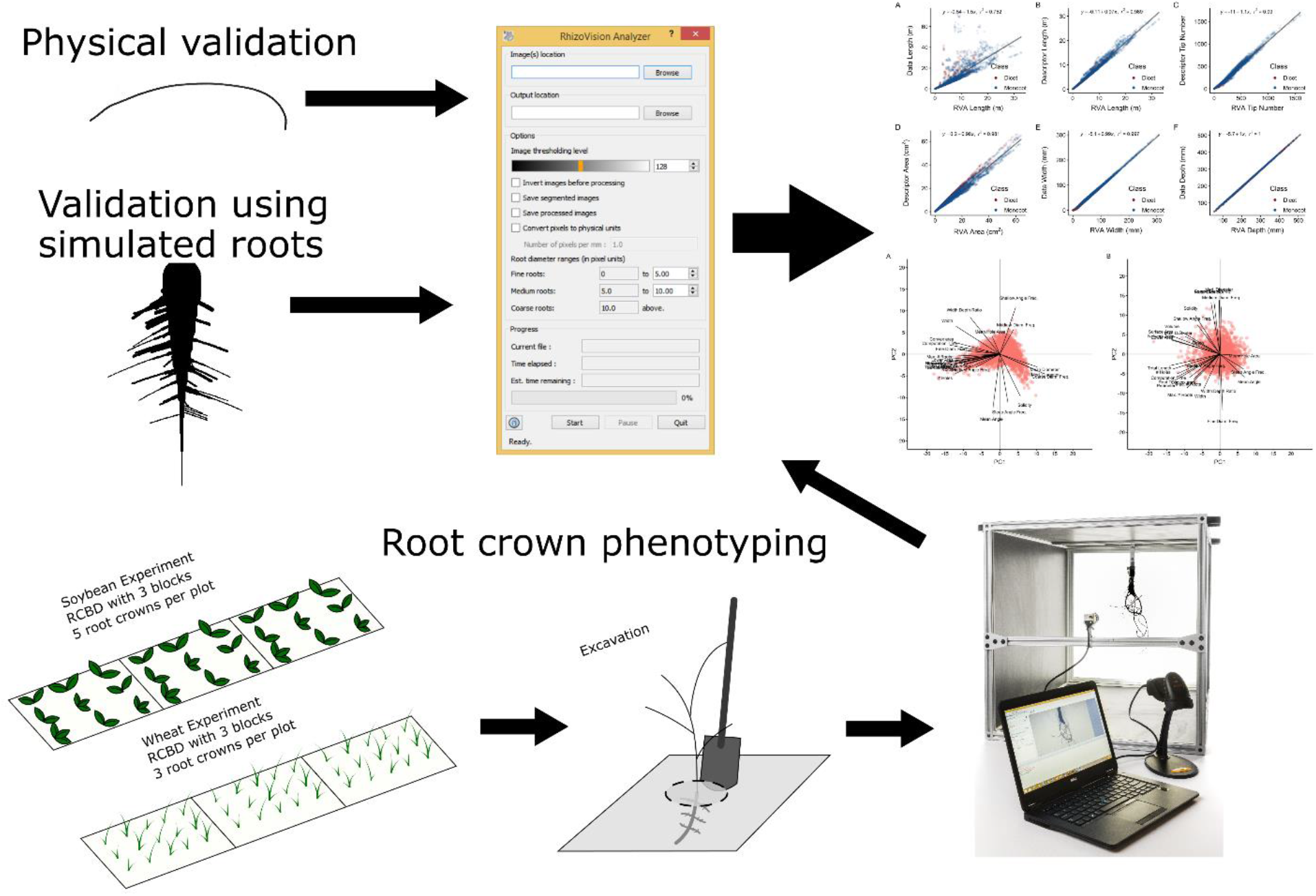
The experimental design of this study to design and validate a new root crown phenotyping platform included constructing a hardware platform, developing a software for imaging, and developing a software for image analysis. The image analysis was validated for physical measures using scans of copper wire and for root measurements using simulated monocot and dicot images. Finally, root crown phenotyping was validated using a soybean experiment in Missouri and a wheat experiment in Oklahoma.

### RhizoVision Crown Hardware

The RhizoVision Crown hardware platform (Fig. 2A) is a backlit solution designed to produce images in which the background is nearly completely white and the foreground (root crown) is nearly black because it is a silhouette. This is achieved by use of a 61 cm x 61 cm LED edge lit flat panel light (Anten, 40 watts, 6000K light color) affixed with epoxy to the back of an imaging box. The imaging box is constructed from T-slotted aluminum profiles (80/20 Inc., Columbia City, IN) that were assembled to generate a box measuring 65.5 cm x 65.5 cm x 91.4 cm. Foamed black PVC panels were custom cut (TAP Plastics, Stockton, CA) and placed between profiles to isolate the interior from outside light. A root crown holder was constructed by attaching a spring clamp to the bottom of a foamed PVC panel measuring 22.86 cm x 30.48 cm. On the top of the root holder panel a screen door handle was attached to assist with the placement and removal of the root holder on the instrument. Detailed images, a schematic plan, and the parts list for the aluminum frame are available as Supplementary Material 1. A root crown is clamped onto the holder, and the holder panel is placed in an indentation designed into the top of the imaging box such that root crowns are consistently placed at the desired position. At one end of the imaging box is the LED panel, and on the other is a CMOS sensor monochrome camera (Basler acA3800-um, Graftek Imaging, Inc., Austin, TX) using a 12 mm focal length lens (Edmund Optics 33-303, Graftek Imaging, Inc., Austin, TX). The camera is connected to a laptop computer USB 3.0 port using a USB 3.0 cable (Micro-B male to A male connectors). For the recommended barcode mode, a USB barcode scanner was also connected to the laptop (Tautronics, Fremont, CA). The imaging software is described in the following section.

**Figure 2.**
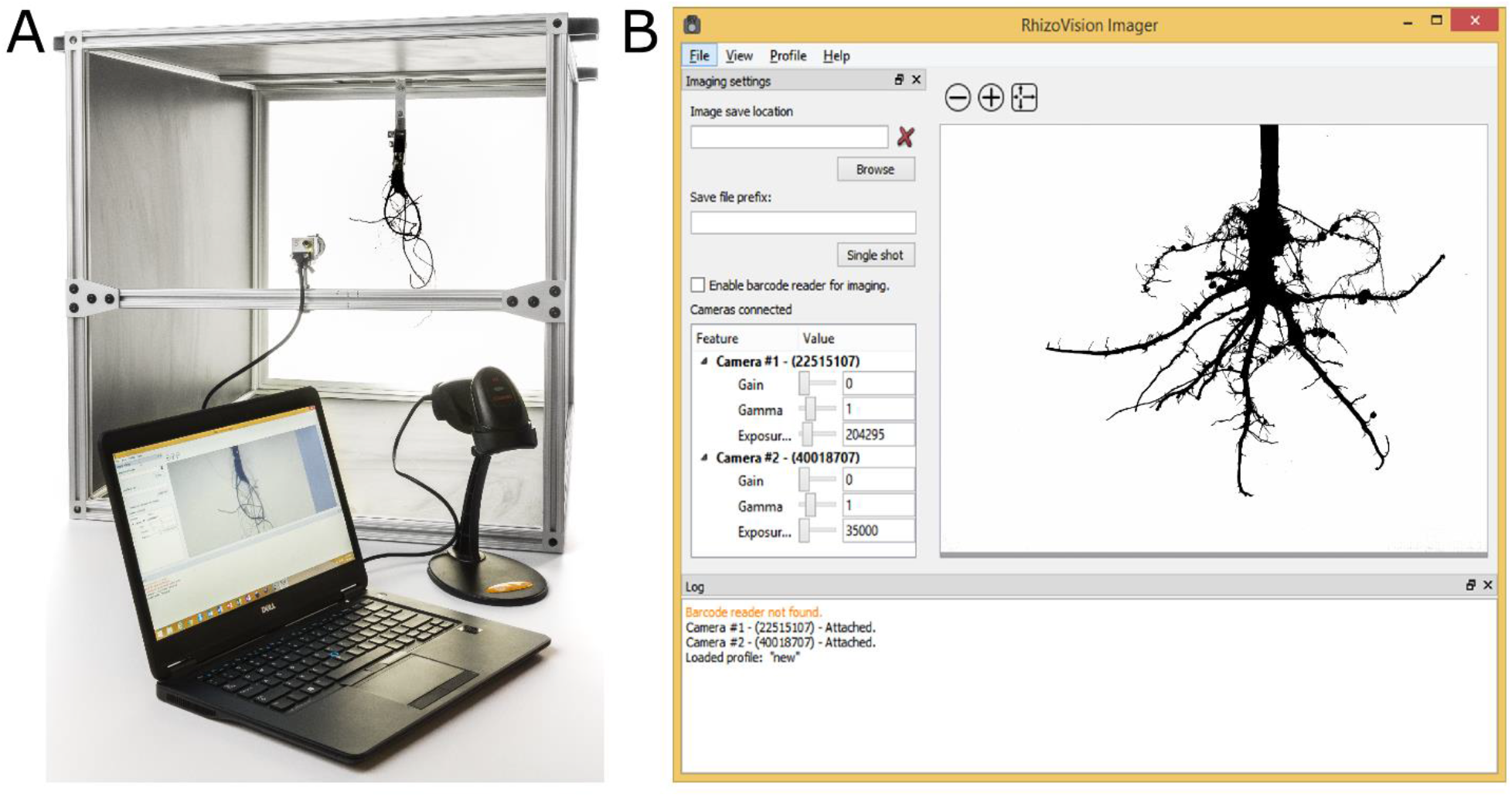
RhizoVison Crown hardware and software for root crown imaging. Root crowns are placed into the imaging unit (A) with a backlit panel for framing the root crown and a laptop connected to a vision camera and USB barcode scanner. The vision camera is controlled using the software RhizoVison Imager (B) which has a user interface for controlling camera settings, provides a live camera view, and image export settings.

### RhizoVision Imager

The imaging hardware is controlled by RhizoVision Imager (Fig. 2B). The software package is open-source and can be downloaded for free from https://doi.org/10.5281/zenodo.2585881 (for x86_64 processor). The program can connect to multiple Basler USB 3.0 cameras and capture images using the Basler Pylon SDK. For each camera, the parameters Gain, Gamma and Exposure Time can be changed to suit the experimental needs (Fig. 2B). Using the lenses mounted on the cameras, the aperture and focus of the lens can be modified.

The program starts with a live view for a connected camera. If multiple cameras are connected, the live view for each can be changed in the View menu. The live view can be zoomed in and out to view a specific area in the image. To start capturing images from the connected cameras, a directory location needs to be specified in which to save the images. For single shots, the user may enter an image file name. File names of all the captured images are appended by the camera number and by the number of times the image was taken with the same name and camera number. This ensures that the images are not overwritten and allows for multiple subsamples of the same biological replicate to be acquired using the same identification.

The program also supports barcode reading for designating filenames and image capture. When a barcode reader is connected to the computer and enabled in Imager, images are captured from all cameras when a barcode is scanned with appended camera number and picture number. The program has a log window, where all the events are logged for review. This includes logging when a new image is captured, camera devices are refreshed or a barcode scanner is attached. The camera settings can be saved as profiles in the program, which may then be reused in later experiments or modified with a text editor. The images can be captured as .BMP, .JPEG, .PNG or .TIFF files. RhizoVision Imager was implemented in C++ using OpenCV, uses the Basler Pylon SDK, and the user interface was developed using Qt, a cross-platform GUI toolkit.

### RhizoVision Analyzer

RhizoVision Analyzer (Fig. 3A) is designed to quickly analyze the images acquired using the RhizoVision Crown platform and the Imager software. Analyzer is open-source and can be downloaded for free from https://doi.org/10.5281/zenodo.2585891 (for x86_64 processors). The overall goal in the design of RhizoVision Analyzer was to create a simple-to-use and robust program that batch processes a folder containing root crown images and outputs a data file with the measures for each sample in a form convenient for data analysis. Analyzer has an option to output segmented images (Fig. 3B) as well as processed images on which visual depictions of the extracted features are drawn on the segmented image (Fig. 3C).

**Figure 3.**
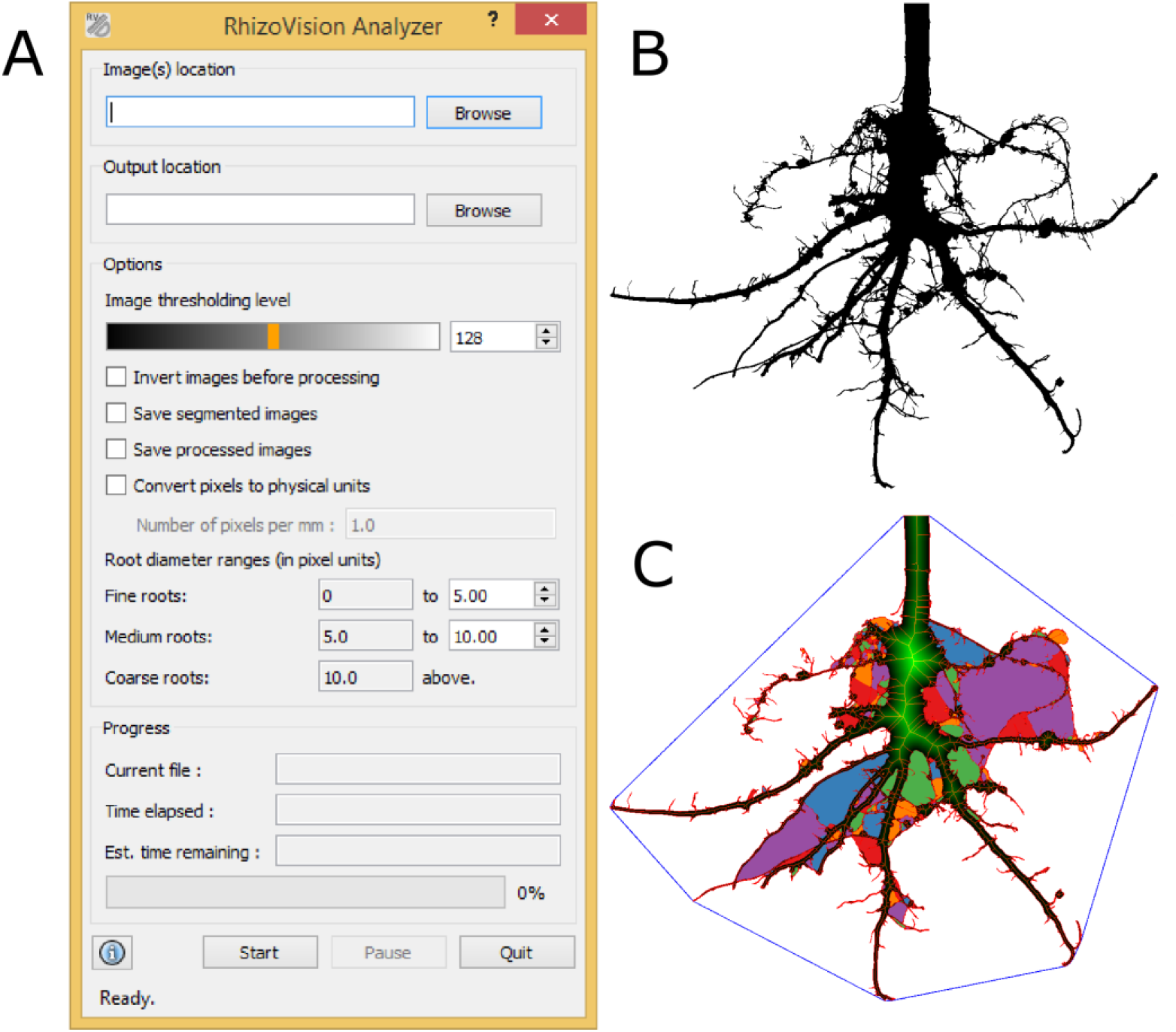
RhizoVision Analyzer for automated batch analysis of root crown images. (A). The software has a user interface (A) for selecting input and output folders, choosing image threshold levels before analysis, classifying root diameter ranges and saving options. The segmented image (B) and feature image (C) are optionally generated by RhizoVision Analyzer. The feature image shows a blue convex polygon that is fit around the entire root system for extraction of Convex Area. The boundary and skeletal pixels are shown in red and the distance transform is shown in green. The “holes” or the background image patches that were disconnected due to the overlapping of foreground pixels are colored for distinction.

Coupled with the optimized image acquisition using the hardware platform, segmentation of the root crown images from the background requires only thresholding of the greyscale values for each pixel with minimal loss of data (Fig. 4B). Thresholded (binary) or greyscale images from other platforms may also be used. The input image may have irregular edges that lead to nonexistent skeletal structures being created (Fig. 4C). Hence, the edges of the input image are smoothed so as to remove irregularities along the edge using the Ramer–Douglas–Peucker algorithm (Ramer, 1972; Douglas and Peucker, 1973). After this procedure, overall shape of a root segment does not change substantially, but the skeletal structure now is simpler and has fewer non-existent lateral roots (Fig. 4D,E,F).

**Figure 4.**
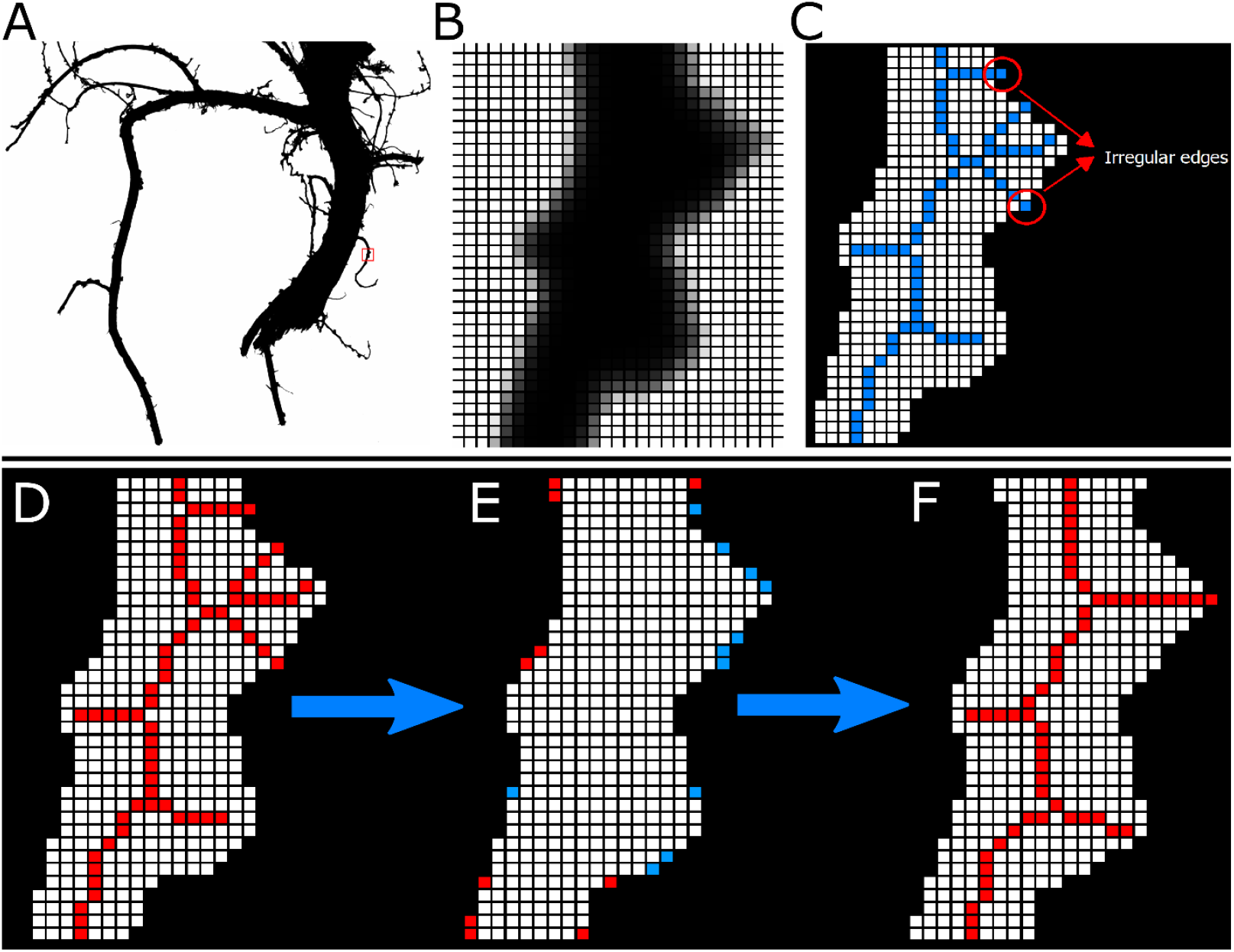
Example of how RhizoVision Analyzer skeletonizes root crown images before extraction of measurements. A small region of interest is selected (A) and magnified (B) for demonstration purposes. The thresholded image of the region of interest shows that due to the irregular edges, the generated skeletal structure contains lateral roots that are non-existent (shown in blue) (C). The skeletal structure of the root is then smoothed to reduce falsely classified lateral roots before line smoothing operation (D). During the line smoothing operation pixels are either added (shown in red) or deleted (shown in blue). Finally, the skeletal structure of the root after line smoothing operation has the falsely classified lateral roots removed (F).

On each row of the segmented and smoothed image, each pixel transition from background to foreground (plant root pixel) is counted, obtaining a plant root count profile along the depth of the root crown, from which Median and Maximum Number of Roots are determined (Fig. 5). Maximum Width and Depth are extracted from this smoothed image. The Network Area of the image is determined by counting the total number of plant root pixels in the image. Further, a convex polygon is fit on the image and the area of this polygon is noted as Convex Area.

**Figure 5.**
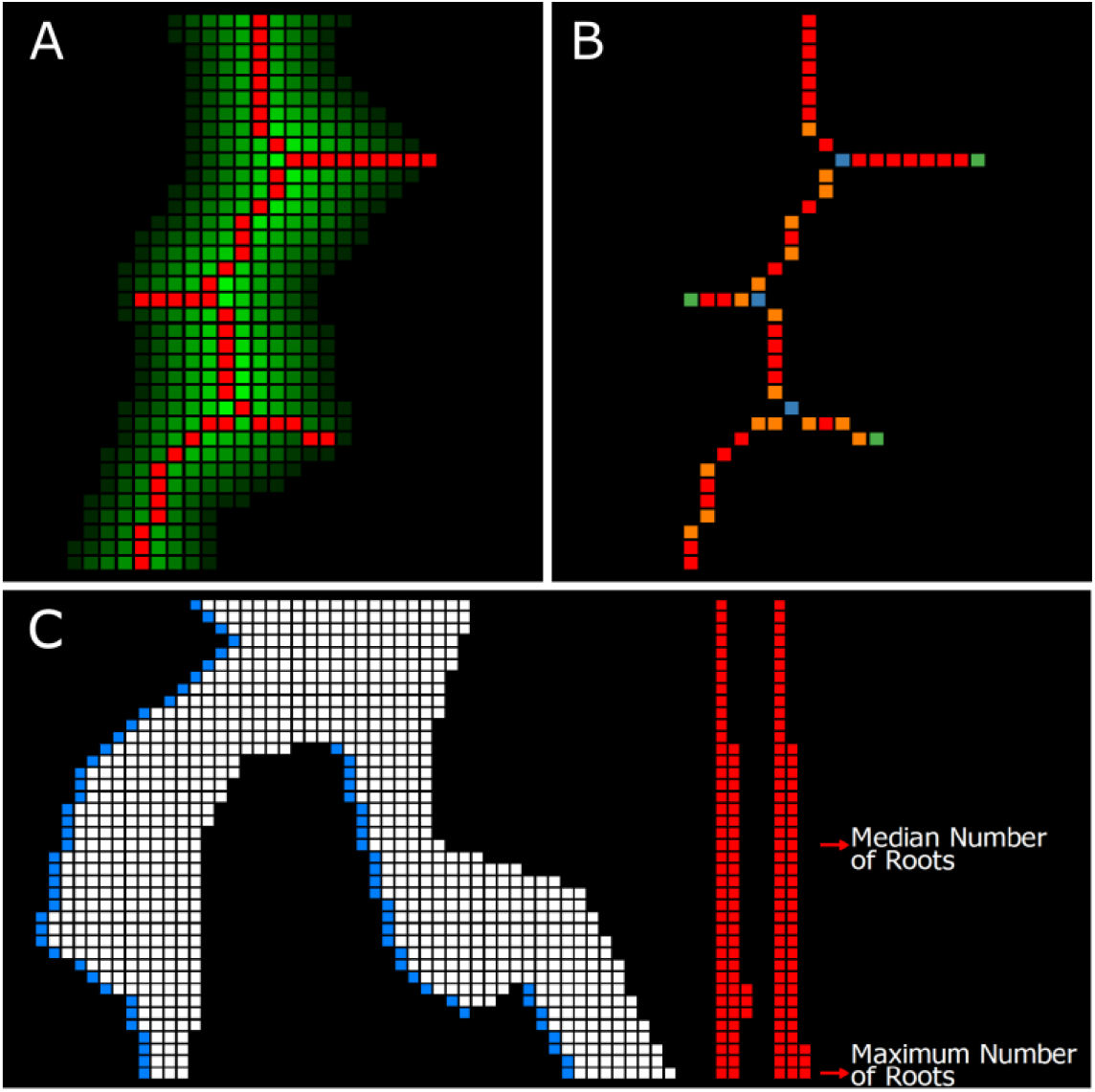
Example of how RhizoVision Analyzer extracts quantitative features from the skeletonized root crown. For each pixel within the root crown skeleton, the corresponding value from the distance map is used to estimate root diameter (A). Topological information is extracted from the skeletal structure such as branch points (shown in blue), root direction change (shown in orange) and end points (shown in green) (B). Finally, for the root counting procedure (C) a pixel transition is marked in a horizontal line scanning operation (shown in blue) for each row and is recorded for counting the number of roots in that row (shown in red).

A precise distance transform is computed on the line smoothed image in order to identify the medial axis. The distance transform (Felzenszwalb and Huttenlocher, 2012) of an image is the map of distance of each pixel to the nearest background pixel. The distance metric used here is the Euclidean distance metric (Fig. 5A). The medial axis is a set of loci on the distance transform that are equidistant from at least two background pixels and is identified from the ridges formed on the distance transform map (Fig. 5A). In order to make a fully connected skeletal structure, additional pixels are added using the connectivity preserving condition from the Guo-Hall thinning algorithm (Guo and Hall, 1989; Lam et al., 1992) and the endpoints of the ridges are connected using the steepest accent algorithm. The contours of the segmented image are identified for determining the perimeter of the plant root image.

Using the generated skeletal structure, topological properties such as the branch points and end points are identified (Fig. 5B). The skeletal pixels connecting one branch point to another branch or end point are identified as root segments. The number of end points are noted as Number of Root Tips. For each skeletal pixel in every root segment, a 40 x 40 neighborhood window is selected. All the skeletal pixels on the root segment of the current skeletal pixel are taken within the window and average angle is computed. Using these angles, the numbers of shallow-angled, medium-angled, and steep-roots in an image are noted as histogram bins, by grouping the computed angles in ranges of 0° to 30°, 30° to 60° and 60° to 90°, respectively. This histogram is normalized and the bins are named as Shallow, Medium and Steep Angle Frequencies. Further, an average of all the angles is computed and noted as Average Root Orientation. A similar normalized histogram is constructed using the skeletal pixels on the root diameter. The histogram bins are allowed for the user to be specified from the user interface of RhizoVision Analyzer. These bins are noted as Fine, Medium and Coarse Diameter Frequency. Also, the Average, Median and Maximum diameters are identified from the diameters of all the skeletal pixels. The plant root area below the pixel having maximum diameter is noted as Lower Root Area. The segmented and edge smoothed image is color inverted and connected component analysis is performed to count the number of Holes and an average of all the sizes of holes is computed to determine the Average Hole Size. Table 1 briefly describes the list of features extracted from the root crown images.

**Table 1.** The list of 27 features extracted from each root crown image by RhizoVision Analyzer.

| Features extracted | Description |
| --- | --- |
| **Median and maximum number of roots** | The number of roots are counted by performing horizontal line scans from left to right in each row through the segmented image. In each of the line scan, we check if there is a pixel value transition from the previous pixel value to the current pixel value on its right side. If the current pixel value changes from 0 to 1, we note that a root is present. The number of roots are recorded from each row of the segmented image and the median and maximum number of roots is determined from these values. |
| **Number of Root Tips** | Computed by counting total number of tip pixels in the skeletonized image. |
| **Total root length** | Computed by counting the total number of pixels in the skeletonized image. |
| **Depth, maximum width and width-to-depth ratio** | The trait values for both depth and maximum width of the root in the segmented image. The ratio of maximum width to depth of the image is noted as width-to-depth ratio. |
| **Network area, convex area and solidity** | Network area is the total number of pixels in the segmented image. The convex hull of a geometric shape is minimal sized convex polygon that can contain the shape. The ratio of network area and the convex area is noted as the solidity. |
| **Perimeter** | Perimeter is the count of total number of pixels in the perimeter image. |
| **Average, median and maximum diameter** | For each pixel on the skeletonized image, the distance to the nearest non-root pixel is computed and using this distance as radius a circle is fitted. The diameter of the circle at each pixel is noted as the diameter at that pixel. We get the list of diameters from all the medial axis pixels and determine the average, median and maximum diameter. |
| **Volume and surface area** | Using the radii determined earlier, the sum of all cross-sectional areas across all the medial axis pixels are noted as volume and the sum of the perimeter across all the medial axis pixels are noted as surface area. |
| **Lower root area** | The lower root area is the area of the segmented image pixels that are located below the location of the medial axis pixel that has the maximum radius. |
| **Holes and Average hole size** | Holes are the disconnected background components and indicative of root branching and complexity. They can be counted by inverting the segmented image. The average hole size (area) is also calculated. |
| **Average Root Orientation** | For every medial axis pixel, the orientation at the pixel is computed by determining the mean orientation of medial axis pixels in a 40x40 pixel locality. The average of all these orientations is noted as average root orientation. |
| **Fine, Medium, Coarse Diameter Frequencies** | From the skeletal image, the medial axis pixels are grouped into fine or coarse roots based on the diameter values at the pixels. |
| **Shallow, Medium, Steep Angle Frequencies** | Given the skeletal image, for every pixel in the medial axis, we get the locations of the medial axis pixels in a 40x40 pixel locality and determine the orientation of these pixels in the locality. This orientation is noted for every medial axis pixel. Given these orientations, we calculate the frequency in bins less than 30, less than 60, and less than 90 degrees. |
| **Computational time** | The time taken to extract traits for every plant root image. |

RhizoVision Analyzer is implemented in C++ using the OpenCV library. The user interface of the program is developed in Qt, a cross-platform GUI toolkit. The program can utilize a CPU’s vectorization facilities using Intel’s AVX 2.0 technology, to execute the algorithms faster on newer computers. All pixel-based measures are converted to appropriate physical units if the user supplies the number of pixels per millimeter before analysis. Depending on the exact computer system, Analyzer can be expected to routinely process each image in a fraction of a second.

### Validation Using Copper Wire and Simulated Root Systems

In order to validate the ability of RhizoVision Analyzer to correctly determine physical measurement units from pixel-based analysis of images, copper wire of different diameter gauges were scanned and analyzed. The gauges used were 10, 16, 22, 28, and 32. Two lengths of wire were used for each gauge for a total of ten. The ground truth diameter was measured using a micrometer. The ten wires were scanned individually at 800 DPI using an Epson Expression 12000XL scanner with a transparency unit using Epson Scan 2 software. These images were processed using RhizoVision Analyzer with 31.496 used to scale from pixels to millimeters.

To validate the diverse root measures generated by the RhizoVision Analyzer software, 10,464 simulated images of dicot and monocot root systems from Lobet et al. (2017) were processed (elapsed time 1 hour 7 mins on an Intel processor with 8-cores, 3.7 GHz of clock frequency, 16 GB of RAM memory). Lobet et al. (2017) define the ground truth data as the known measurements from the three-dimensional simulations and the descriptor data as those derived from projected 2D images using RIA-J image analysis.

### Field Sites and Root Crown Phenotyping

#### Phenotyping soybeans in Missouri

A F5-derived soybean recombinant inbred line (RIL) population derived from a cross of PI 398823 × PI567758 was planted at the Bradford Research Center near Columbia, MO on a Mexico silt loam soil (fine, smectitic, mesic Aeric Vertic Epiaqualf). The parental lines of this population were previously identified to differ in top-soil root architecture based on the characterization of soybean diversity panel (F.B. Fritschi, unpublished). Pre-plant soil tests indicated that no P or K fertilizer application was necessary. Prior to sowing, the seedbed was prepared by one pass with a disc to approximately 0.15 m depth, which was followed by a pass with a harrow. The 185 RILs and two parental lines were sown in a randomized complete block design with three replications on 14 May 2017 at a density of 344,000 plants ha-1 in single 3 m long rows with a row spacing of 0.76 m. Weed control consisted of a pre-plant burn-down application of glyphosate (0.73 kg ha-1 a.i.) and post planting applications of acetochlor (0.6 kg ha-1 a.i.), bentazone (0.27 kg ha-1 a.i.), and clethodim (0.03 kg ha-1 a.i.), and these herbicide applications were supplemented by manual weeding as needed. Additionally, two applications of zeta-cypermethrin S-cyano (0.1 kg ha-1 a.i.) were conducted to control insects.

Five root crowns for each plot were excavated at the beginning of the R6 stage the week of 21 September 2017 using a shovel. For each focal plant, the shovel was inserted such that the width of the blade was parallel to the row and mid-way between two rows on each side of the plant. The blade was inserted as deeply as possible and on the second insertion the shovel was leveraged in order to pry up the plant. The soil was very loose root crowns only needed shaken to remove the majority of soil and were not washed. The root crowns were imaged using RhizoVision Crown in the field using a gasoline electric generator for power. The lens of the camera was placed at a working distance of 56 cm from the center of the root crown (the bottom of the clamp) for a resolution of 12.7787 mm per pixel. The lens aperture was set to f/11.0 to maximize the depth of field to accommodate the 3D root crown. Exposure time was set to 14 ms and gamma was set 3.9 in order to optimize contrast. Roots were placed in the orientation that appeared as the widest to the user in order to standardize measurements.

#### Phenotyping wheat in Oklahoma

The wheat population is a recombinant inbred line (RIL) population with 184 F_5:7_ lines derived from a cross between TAM 111 × TX05A001822. The population was created for mapping QTL or genes contributing to a number of important agronomic traits. ‘TAM 111’ is one of the most planted hard red winter wheat cultivars in the Southern High Plains and has adapted to both dryland and irrigated conditions (Lazar et al., 2004) while TX05A001822 is an advanced breeding line with superior bread making quality from the Texas A&M AgriLife Research.

The population was planted in a randomized complete block design with three replications of 1.5 m wide by 0.9 m long plots with seven rows and seeded at a rate of 148 kg ha^−1^ on 11 November 2017 at Burneyville, Oklahoma. The field was clean tilled prior to planting and rain-fed with no supplemental irrigation. Fertilization was first pre-plant incorporated with 56 kg ha^−1^ nitrogen and then top-dressed with 56 kg ha^−1^ nitrogen on 23 January 2018 based upon rainfall. Phosphorous and potassium concentrations were sufficient based on soil test results prior to planting. Weeds were controlled with 247 kg ha^−11^ of glyphosate at planting and 0.02 kg ha^−11^ of Glean XP at Zadoks growth stage 13. Post-emergence application of 1.12 kg ha^−11^of 2,4-D was used on 14 February 2018 for broadleaf weed control.

Root crowns were excavated near grain maturity on 14 – 15 May 2018. Several plants were harvested with a single excavation because of the high population density. The shovel was inserted parallel to the row with its back against the neighboring rows on each side of the focal plants and a whole group of plants was lifted out then placed into a large plastic bag with a barcode label affixed for sample identification. These bags were taken to the washing station where the group of root crowns in soil were placed in water with dish soap and allowed to soak in one of 20 plastic bins. After soaking and gently moving back and forth in water to remove most soil, the root crowns were removed and washed with a water hose spray nozzle with light pressure for a few seconds to clean more thoroughly. The group of plants remained together and were placed back into the plastic bags. These bags were transported back to the lab and kept in a cold room for one week while imaging using RhizoVision Crown. Three plants were selected from the group and the barcodes were used for triggering image acquisition and saving file names. The lens of the camera was placed at a working distance of 51.5 cm from the center of the root crown (the bottom of the clamp) for a resolution of 14.0315 mm per pixel. The lens aperture was adjusted to f/11.0. Exposure time and gamma were set to 14 ms and 3.9, respectively.

### Statistical Analysis

Statistical analyses were employed by using R version 3.5.1 (R Core Team, 2018) through RStudio version 1.1.45 (RStudio, 2016). Linear regressions were fit using the ‘lm’ function. Principal component analysis was conducted using the ‘prcomp’ function after scaling and centering the data. The R package ‘reshape2’ (Wickham, 2007)was used to format the data before plotting. The R package ‘ggplot2’ (Wickham, 2016) was used for data visualization. Other packages used included ‘dplyr,’ ‘purr,’ and ‘patchwork.’ Broad-sense heritability was calculated based on (Falconer and Mackay, 1996) as:

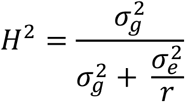

The variables 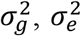 and *r* represent the variance of the genotype effect, variance of the environment effect, and the number of replicates, respectively. The variances were obtained by fitting a mixed model including genotype as a random effect and replicate as a fixed effect using the lme4 package (Bates et al., 2015). The data for the five root crowns of soybean and three root crowns of wheat from each plot were averaged before subsequent analysis.

## RESULTS

### An integrated hardware and software platform for accelerated phenotyping and knowledge generation

The RhizoVision Crown hardware and software platform builds upon previous developments for root crown phenotyping in order to increase usability and reliability. Specifically, the goals were to optimize the stages of sample loading, recording sample identification, image acquisition, image analysis, and data analysis. At the same time, special consideration was given to making sure the platform could be used by as many researchers as possible by developing open hardware that can be built by most organizations and free software that is ready-to-run on the widely available Windows operating system with little technical knowledge.

In order to ensure reproducible imaging of crop root crowns that would allow 100% success rates during image analysis, a backlit solution was chosen (Fig. 2A). An LED flat panel is mounted behind the root crown and a monochromatic machine vision camera faces the light panel and is focused on the root crown. Root crowns are loaded by attaching them to a clip that is affixed to a board that serves as a lid for the opening through which the root crowns are inserted into the imaging box. This board fits an indentation on top of the instrument as to ensure root crowns are loaded into consistent positions, and a handle allows for easy manipulation of the board and replacing of the root crowns. This setup ensures that all images acquired have a white background, with the captured root crown silhouette being primarily black and dark grey (greyscale).

The camera is attached to a laptop computer with a USB cable that supplies power to the camera, sends commands from the laptop to the camera, and transfers image data from the camera to the laptop. The RhizoVision Imager software (Fig. 2B) connects to the camera and provides a live view, allows modifying camera settings, saves setting profiles, and acquires images. Sample identifications (file names) can be typed in for single shots, or a barcode reader can be used for greater throughput and accurate tracking of sample identity. The barcode setting allows image acquisition to be triggered after a barcode label is scanned and saves the resulting image with the barcode string as the file name. After image acquisition, the root crown is replaced and the process repeated, with throughputs achievable of at least 6 root crowns per minute if previously excavated and cleaned. The dependencies of the Imager software are use of Basler machine vision cameras and installation of the freely available Pylon runtime from the camera manufacturer. All RhizoVision software described are open-source with a modified GPL license, designed for Windows 10, and do not require installation.

Once images are acquired, a separate software named RhizoVision Analyzer (Fig. 3A) is used for extraction of phenes from the images in batch mode. The user simply provides the directory containing the images, an output directory for generated data, and a greyscale thresholding level for segmenting roots from the background before pressing “Start”. Additional options include saving segmented images and feature images that overlay the features on the segmented image. The pixel units can be converted to physical units if the user supplies the pixels per millimeter. Finally, the diameter ranges for fine, medium, and coarse roots can be defined by the user. The output directory includes a data file with a column for the sample names followed by the 27 extracted measurement columns, and a separate metadata file that stores the user-defined options. The Analyzer software has no additional dependencies to run.

The hardware platform optimizes image acquisition of root crowns to increase throughput and ensure successful image processing. A high level of image quality is achieved with approximately $1,200 USD of hardware that can be assembled by most laboratories, including the aluminum profiles, plastic panels, LED panel, camera, and lens but excluding a computer. The RhizoVision software is free, open-source, and can be used independently of the imaging box assembly. The Imager software coupled with the imaging box assembly allows high contrast root crown images to be generated with relative ease and speed. The Analyzer software can process each image in a fraction of a second and the data output is in a format ready for data analysis pipelines. This integrated platform could contribute to root biology by allowing more labs to conduct root crown studies on diverse topics and could serve as a benchmark for other integrated hardware and software platforms (Lee et al., 2018).

### Physical calibration

In order to ensure that the correct physical units were generated by the RhizoVision Analyzer software, copper wires of known diameters ranging from 0.2 – 2.57 mm were scanned with a flatbed scanner at 800 DPI and the correct pixels per mm conversion was supplied to Analyzer. Regression of the computed diameters versus caliper-measured diameters showed nearly exact correspondence (y = 0 + 1 x, R^2^ = 0.99, p < .01), which indicates the physical units provided by Analyzer are accurate when the user supplies the correct pixels to mm conversion (Fig. 6).

**Figure 6.**
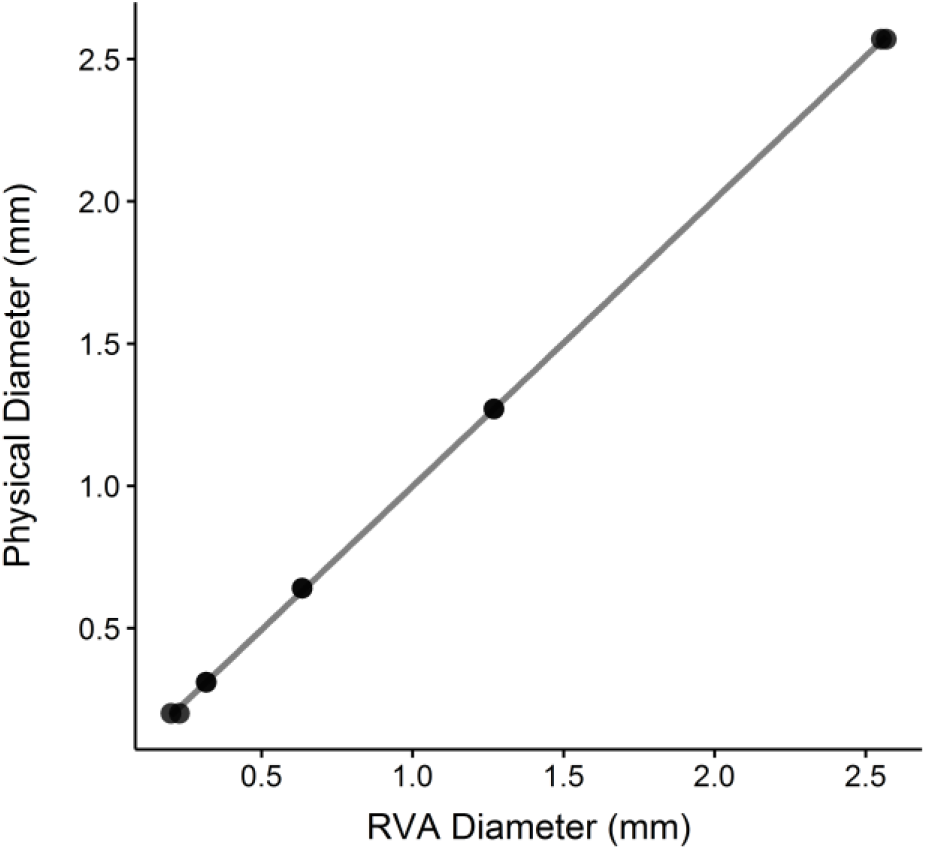
Regression between diameters of copper wires extracted using RhizoVision Analyzer (RVA) and caliper-measured diameters (each diameter has two points).

### Validation using simulated root system images

In order to compare the root measures from RhizoVision Analyzer to ground truth data and data derived from root images using other software, more than 10,000 simulated root images were analyzed and features found in common were compared from Lobet et al. (2017). The ground truth total root length was under-estimated by Analyzer (y = −.54 + 1.5 x, R^2^ = 0.75, p < .01) (Fig. 7A), which is to be expected as the original simulated roots were three-dimensional but the processed images are projected to two dimensions. The descriptor length provided was similar to the Analyzer length (y = −.11 + 0.97 x, R^2^ = 0.99, p < .01) (Fig. 7B), indicating that Analyzer performs similarly to the previously-used software. Tip number (y = −11 + 1.1 x, R^2^ = 0.99, p < .01) (Fig. 4C), root crown area (y = .2 + 0.96 x, R^2^ = 0.98, p < .01) (Fig. 7D), root crown maximum width (y = −5.1 + 0.99 x, R^2^ = 0.99, p < .01) (Fig. 7E), and root crown maximum depth (y = −5.7 + 1 x, R^2^ = 1, p < .01) (Fig. 7F) all indicate that Analyzer extracts phenes that have the same physical units (slopes equal one) and strong correlations with the ground truth and with the phenes extracted from other software.

**Figure 7.**
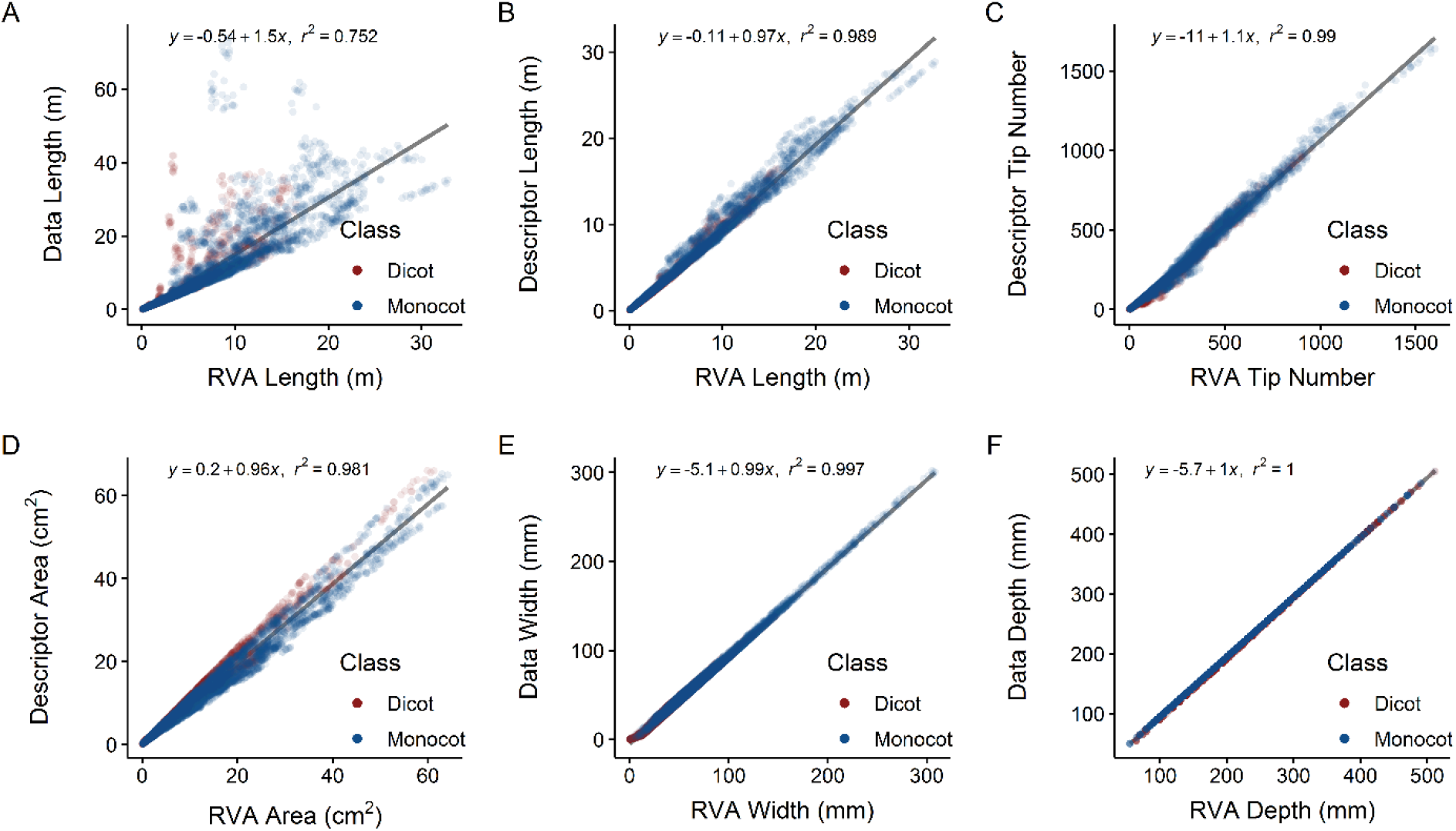
Correlations and linear fit equations between root features extracted using RhizoVision Analyzer (RVA) and ground-truth simulated root data or published original descriptors. Scatter plots include linear regressions of (A) RVA length against ground-truth data length, (B) RVA length against descriptor length, (C) RVA tip number against descriptor tip number, (D) RVA area against descriptor area, (E) RVA width against descriptor width, and (F) RVA depth against descriptor depth.

### Phenotyping soybean and wheat root crowns

In order to validate the entire hardware and software platform, 2,799 of soybean root crowns of 187 lines grown in Missouri and 1,753 images were acquired of wheat root crowns of 186 lines grown in Oklahoma. In Missouri, images were acquired in the field with the imaging system powered by a gasoline generator on the same day the root crowns were excavated. In Oklahoma, images were acquired after wheat root crowns were brought to the lab, stored in a cold room, and imaged within two weeks. In both cases 100% of the root crowns were imaged using RhizoVision Imager and successfully processed by RhizoVision Analyzer, indicating the hardware provides reproducible images that are optimized for image analysis irrespective of plant species. On a computer with an 8-core Intel processor with 16GB of RAM, analysis of the 2,799 soybean images took 17 minutes and the 1,753 wheat images took 11 minutes.

The means and standard deviations were computed for the extracted phenes (defined in Table 1) independently for the wheat and soybean populations (Fig. 8) grown at the two different sites. The average total root length of soybean root crowns was 1.7 ± 1.3 m, number of root tips was 368 ± 264, maximum width was 123 ± 55 mm, and the depth of the roots was 127 ± 30 mm. In general, the entire root crown fit within the field of view of the camera so width and depth measurements are accurate. The soybean root crowns showed solidity values of 0.21 ± 0.09, the median root diameter of 1.4 ± 0.7 mm, hole number of 119 ± 164 and average hole size of 7.5 ± 9.5 mm^2^. Finally, the average root orientation of every pixel in the skeletal structure of the soybean root was 42.5° + 2.9° from horizontal. The average total root length of wheat root crowns was 3.2 ± 1 m, number of root tips was 606 ± 205, maximum width was 78.5 ± 19 mm, depth of the roots was 152 ± 29 mm, solidity was 0.29 ± 0.08, median root diameter was 0.8 ± 0.2 mm, hole number was 499 ± 240, hole size was 3.2 ± 1.9 mm^2^, and the average root orientation of every pixel in the skeletal structure of the wheat root was 49.2° + 2° from horizontal.

**Figure 8.**
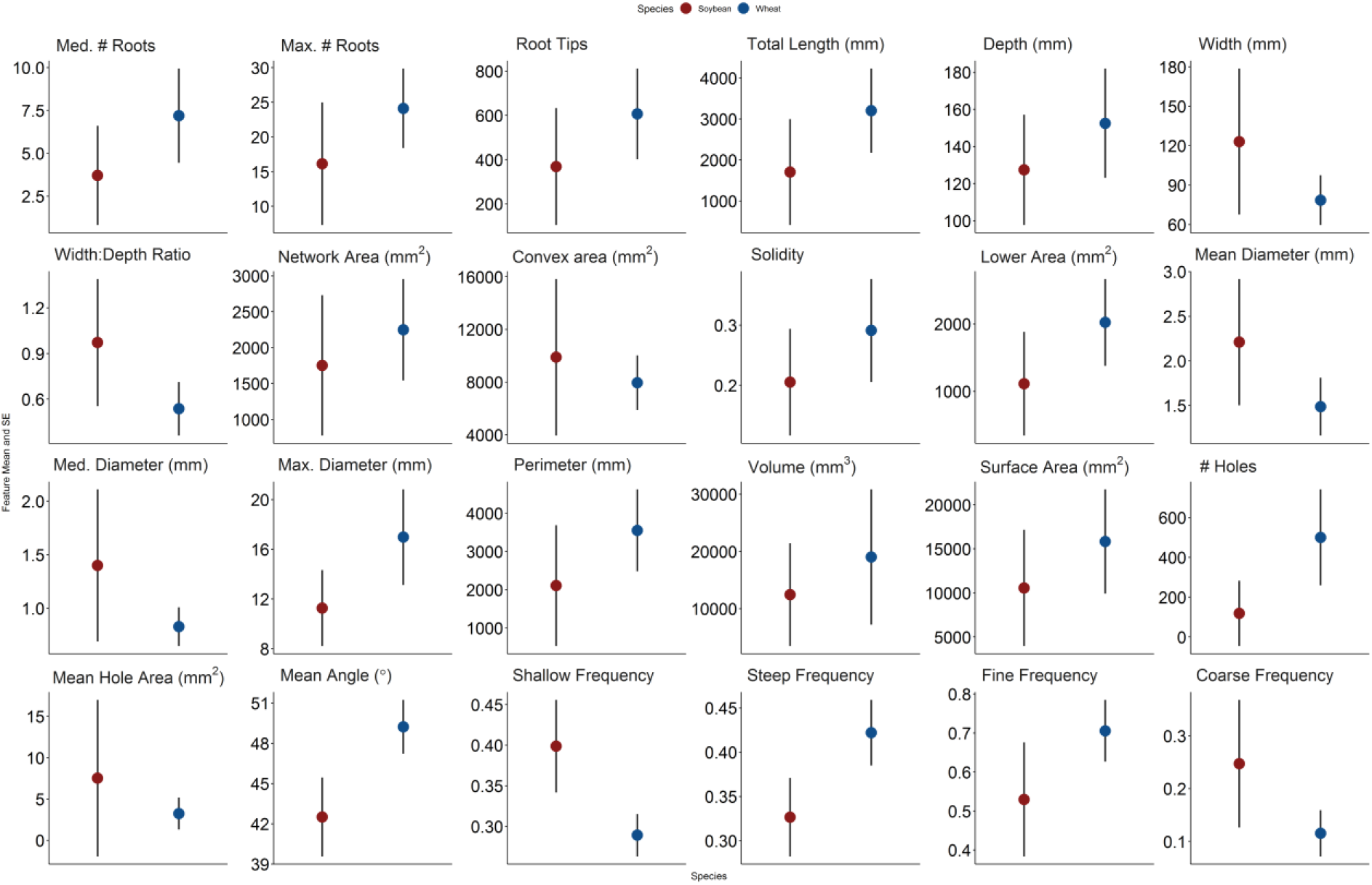
Summary of means and standard errors of various features extracted from soybean (n = 2,799) and wheat (n = 1,753) root crown images using the RhizoVision Crown platform.

Principal component analysis was used to identify the major linear phene combinations that maximize the multivariate variation (Fig. 9A). Principal components (PC) 1 and 2 explained 51.9% and 13% of the multivariate variation, respectively, for the phenes extracted for soybean root crowns. The phenes that loaded most strongly onto PC 1 were size-related phenes such as total root length, perimeter, number of root tips, number of holes, several measures of root areas, and some contribution from diameter measures. PC 2 was dominated by the mean angle and angle frequencies. PCA analysis of wheat root crowns (Fig. 9B) showed that the PC 1 and 2 explained 35% and 27% of the multivariate variation, respectively. The phenes that loaded onto PC 1 were size-related phenes such as total root length, perimeter, number of root tips, number of holes, and maximum diameter. PC 2 was strongly dominated by median diameter and the diameter frequencies.

**Figure 9.**
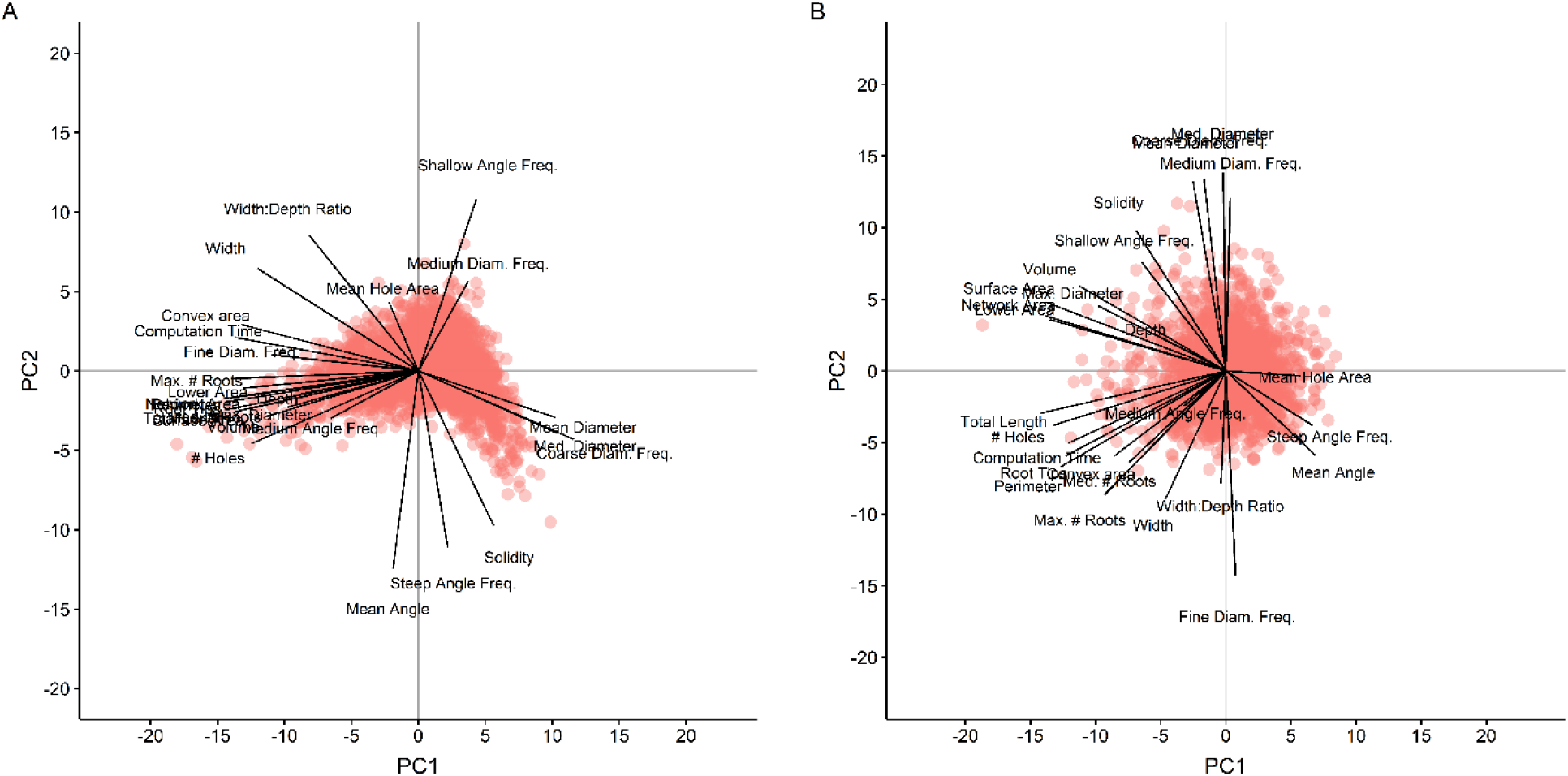
Principal component analysis of root crown features from the soybean (A) and wheat (B) datasets. Points represent the scores of principal components 1 and 2 (PC1 and PC2) for each species. Labelled lines demonstrate the correlation of feature values to principal component scores.

In order to evaluate the possibility to use these root phenes for breeding, broad-sense heritabilities were computed for the phenes extracted from soybean root crown images (Fig 10A). A majority of the phenes had heritabilities greater than 0.5. The maximum heritability was observed with maximum number of roots at 0.74. The phenes with lower heritabilities were the ratios, mean angle and the orientation frequencies. Heritabilities for the wheat root crowns were generally lower, ranging from 0 to 0.22 for maximum width (Fig. 10B).

**Figure 10.**
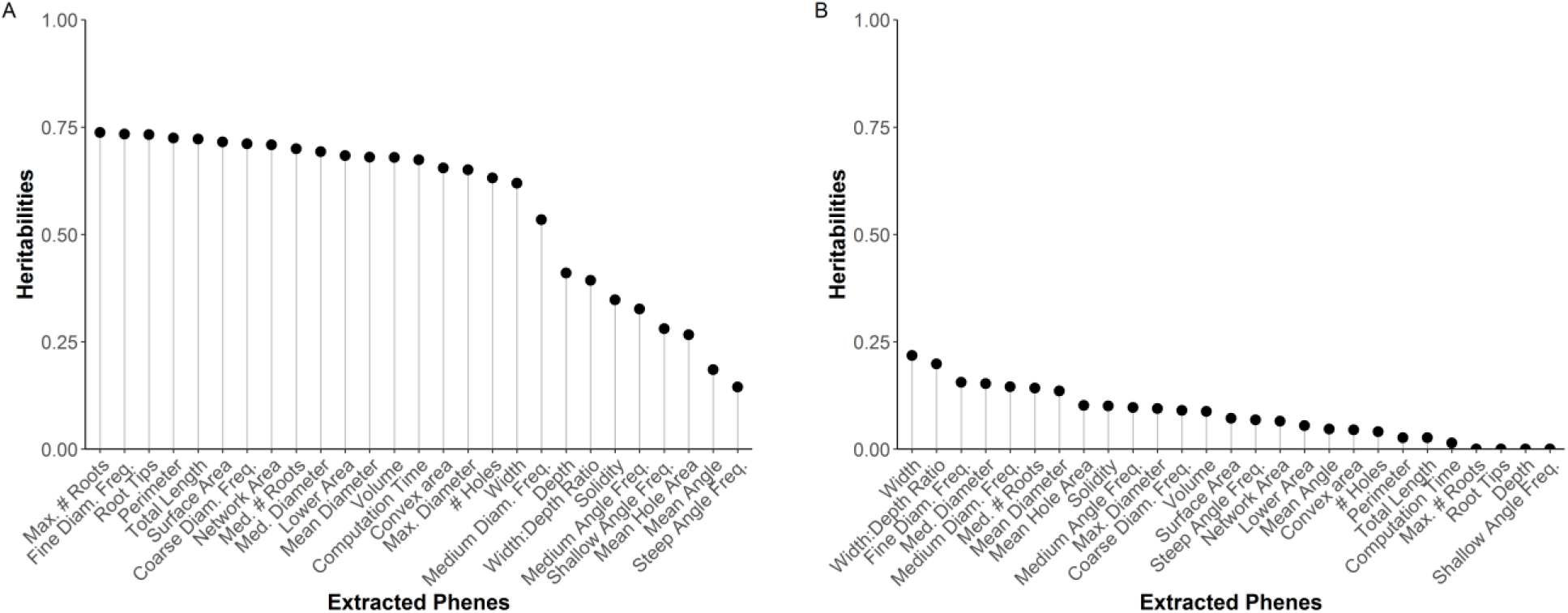
Heritabilities of each phene extracted using RhizoVision Analyzer for soybean (A) and wheat (B) datasets.

## DISCUSSION

Over the past few years, the throughput, reliability, and standardization of root crown phenotyping has been increased using digital imaging and image-based analysis software, such as DIRT (Bucksch et al., 2014), REST (Colombi et al., 2015), and M-PIP (Seethepalli et al., 2018). However, increasing hardware-software integration specifically for root crown phenotyping is promising to further increase throughput and reliability. Minimizing cost, increasing throughput, and improving reliability are key demands for developing high-throughput root phenotyping platforms. The integration of the RhizoVision Imager software with the RhizoVision Crown hardware platform facilitates phenotyping with the end-user in mind by utilizing a backlit approach to capture easily segmentable images and a simple clip-and-replace system for replacing root crowns after imaging. RhizoVision Imager allows live view so that the user may verify that the images are high-contrast and framed correctly, stores camera settings, and has a barcode scanning mode that saves images with the sample identification. The improved quality of images captured enables greater accuracy and precision of root crown measurements, and simultaneously broadens the metrics used to characterize roots (Topp et al., 2016). The ergonomics evident in the hardware and control software facilitate high-throughput image acquisition.

RhizoVision Imager and RhizoVision Analyzer are designed to be used by any user in any lab. Both provide a graphical interface that is intuitive for new users, can be installed by simply downloading a binary archive to a local directory, and eliminate the need for uploading large files to the cloud before feature extraction like DIRT (Das et al., 2015). RhizoVision Analyzer was extensively validated with copper wires of known lengths and diameters as well as with 10,464 simulated images of dicot and monocot root systems with no errors or failures. Excellent agreements were observed between root phenes like length, tip number, root crown area, root crown maximum width and root crown maximum depth extracted using Analyzer and published data of the simulated images. Furthermore, the platform was validated with a phenotypic screen of field-excavated root crowns from soybean and wheat populations.

The soybean and wheat experiments occurred at different times and at different sites so a direct statistical comparison is not possible. However, in both experiments root crown phenotyping occurred after flowering so root crowns were mature. Therefore, the differences observed between the species may be representative of the intrinsic differences. For example, the mean and median root diameters of root crowns are smaller for wheat compared to soybean as would be expected. Wheat root crowns are also typically less wide and with steeper angles due to the growth of nodal roots as opposed to the shallow angles of first order laterals in soybean. While the heritabilities of features for soybean were typically greater than 0.5 with a maximum of 0.74, the maximum observed for wheat was only 0.22. Possibly this indicates that intrinsic differences between the species make the wheat root crown less suitable for phenotyping using this method. For example, the smaller diameter wheat roots are more flexible and when suspended orient downwards, and so differences among genotypes may be obscured. However, root crown phenotyping of field-excavated wheat root crowns was previously used to confirm shallow and steep angles of lines measured in a lab-based seedling screen with success (Maccaferri *et al*., 2016), which indicates the lower heritabilities observed here may not be due to an inherent incompatibility of the method. Another explanation for low heritability is simply that there is not substantial genetic variation for these root phenes present in the RIL population used which is possible because the parents were not selected based on root characteristics, while the soybean parents were selected based on contrasting root system architecture. For species with more flexible roots, refinements to the protocol such as laying the root crown on a flat surface rather than suspending and including more sup-replicates from each plot should be investigated. The imaging box described here could easily be oriented to have the backlight facing up for this use. Additional image-based measures could further improve plant classification and characterization of root topology, for example extracting new root phenes such as lateral root branching density or angles and lengths of specific classes of roots through optimized algorithms. Incorporation of morphometric descriptors (Bucksch et al., 2017) could simplify representation of data, such as persistent homology (Li et al., 2018).

In conclusion, RhizoVision Crown is a cost-effective and high-throughput platform that has the potential to increase access to technologies for root crown phenotyping. The platform builds upon previous platforms (Grift et al., 2011; Bucksch et al., 2014; Colombi et al., 2015; Seethepalli et al., 2018) by optimizing image acquisition using a backlight and the barcode option, using custom imaging software designed for phenotyping, and use of image analysis software with a simple graphical interface designed for batch processing. All software are free and ready-to-use on Windows 10. The platform has been validated using ground-truth measures of a simulated dataset and successfully extracted root phenes from field-excavated root crowns of a cereal and a legume species. The ergonomics of use, the integration of all hardware and software, and the extensive validation tests serve as a benchmark for other plant phenotyping platforms. This technology will increase access to root crown phenotyping as a method to acquire data for functional phenomics (York, 2019), genetic mapping, use in breeding programs, and understanding how root phenes can address agricultural unsustainability and food insecurity.

## Supporting information

Supplemental hardware plans and images

## ACKNOWLEDGMENTS

**General:** We thank Frank Maulana, Bryce Walker, Wangqi Wang, Tadele Kumssa, Jarron Peoples, Franco Guadarrama, Willie Hart, Matt Hogan, Erika Phillips Cheng Lin Chai, and Erica Judd for root excavation, washing, and imaging of the wheat population.

## Author contributions

The RhizoVision Crown platform was conceived by L.M.Y. The software were developed by A.S., with contributions to algorithm development from A.Z. The hardware and software had input from all authors throughout development. H.A. managed the soybean field experiments and conducted root crown phenotyping with F.B.F. in Missouri using seed from Z.L. Wheat experiments were conducted by X.F.M., E.B.B., and X.L. in Oklahoma using seed from S.L., while H.G. organized wheat root crown phenotyping. A.S., H.G., L.M.Y., M.G. and X.L. wrote the first draft of the manuscript and all authors gave input and approved the final version.

## Data availability

The wire and root crown image sets, tabular data, and R code for statistics and graphing are available online: http://doi.org/10.5281/zenodo.3380473. The simulated root images are available as described in Lobet et al. (2017) at: http://doi.org/10.5281/zenodo.208214. RhizoVision Imager is available at: http://doi.org/10.5281/zenodo.2585882. RhizoVision Analyzer is available at: http://doi.org/10.5281/zenodo.2585892.

## SUPPLEMENTARY MATERIAL

Supplementary plans S1 is included as PDF with images of the completed hardware platform, schematic drawings, and a parts list for the aluminum structure.

