## Supplemental hardware plans and images for "RhizoVision Crown: An Integrated Hardware and Software Platform for Root Crown Phenotyping"

### Hardware plans, images, and details (plans and parts last 2 pages)

A

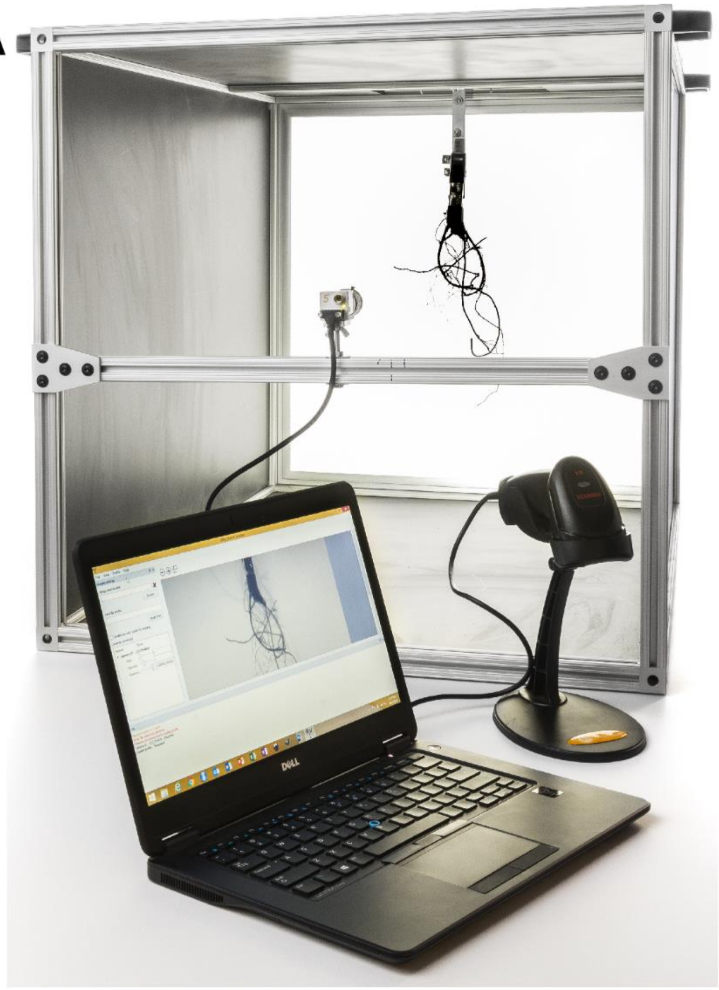

B

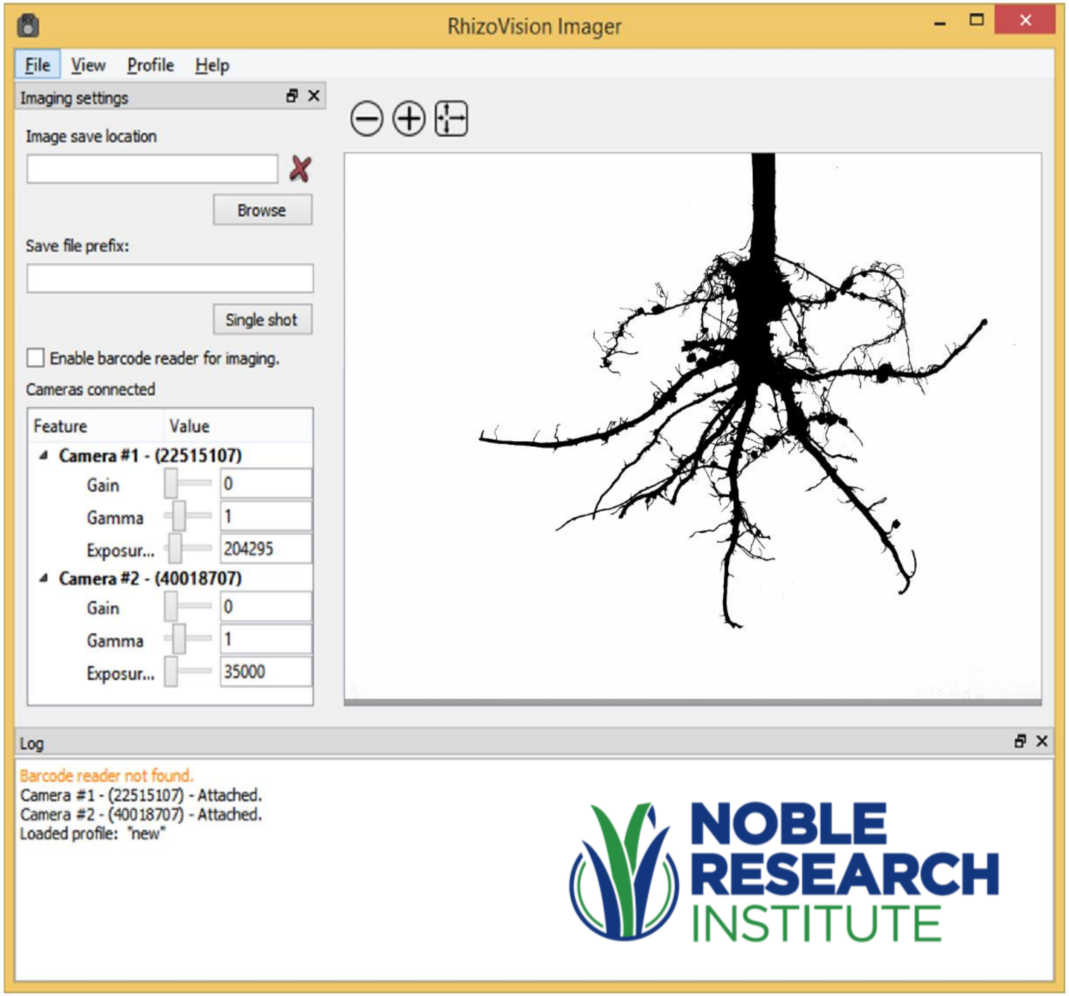

7-hole plates on the back were used to seat light panel then adhesive was added around the interior to keep it in place. Added dimmer but not useful.

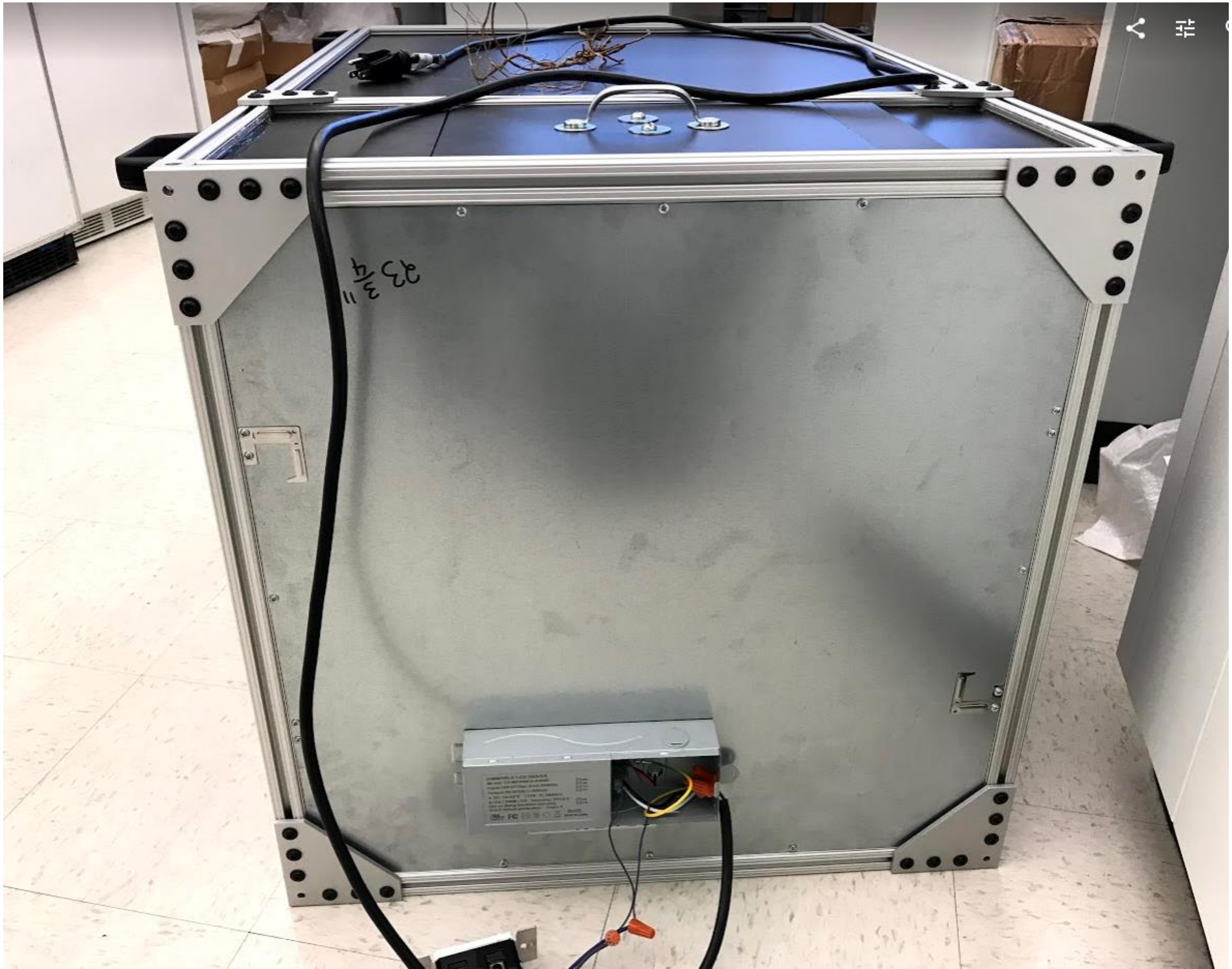

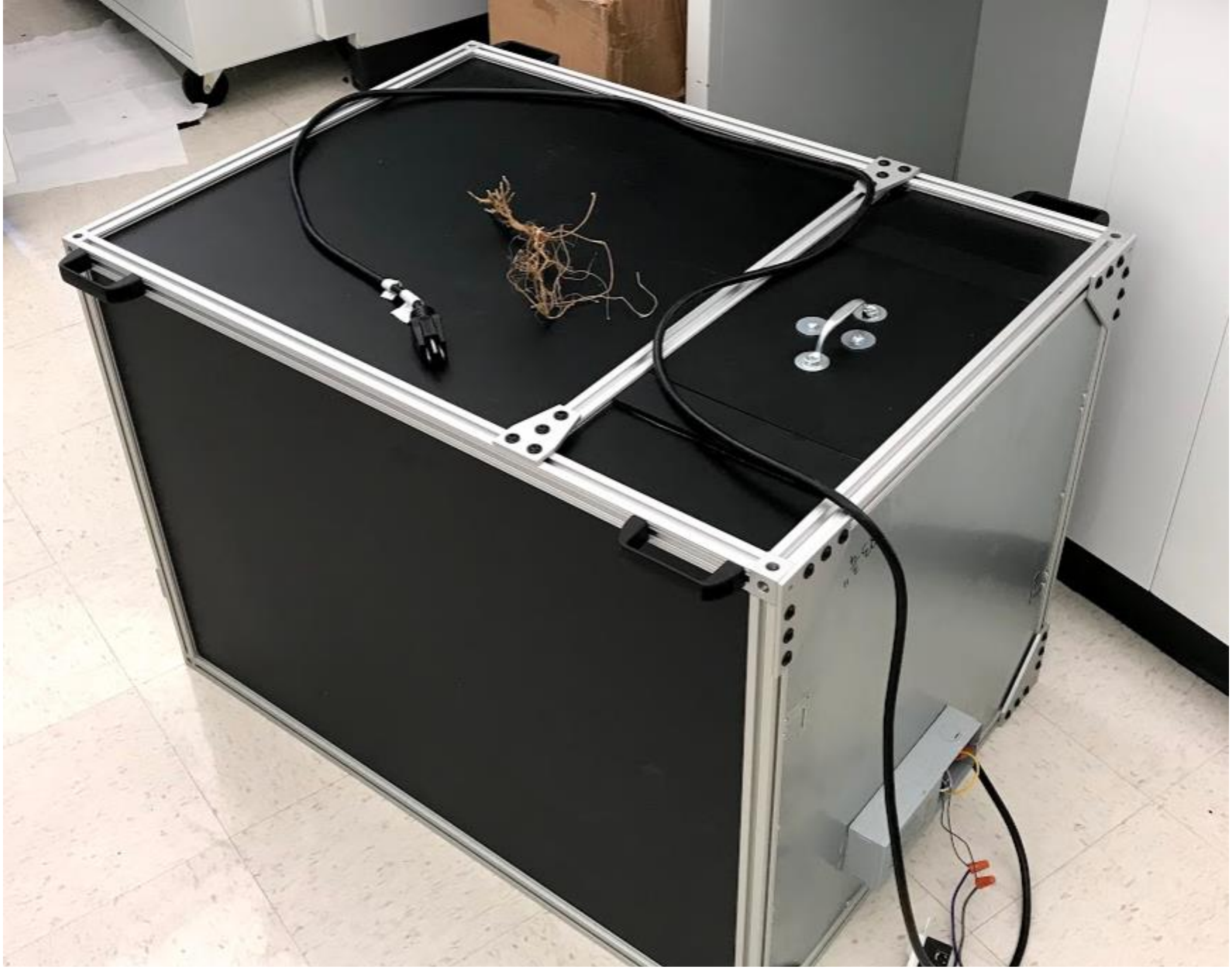

Note this version used corner connectors rather than end-fasteners. The included plans are for use of end fasteners to save money.

Here you can see two ways to mount the camera (ask for the ¼ 20 adapter for the Basler camera to mount on tripod post), either in portrait or landscape.

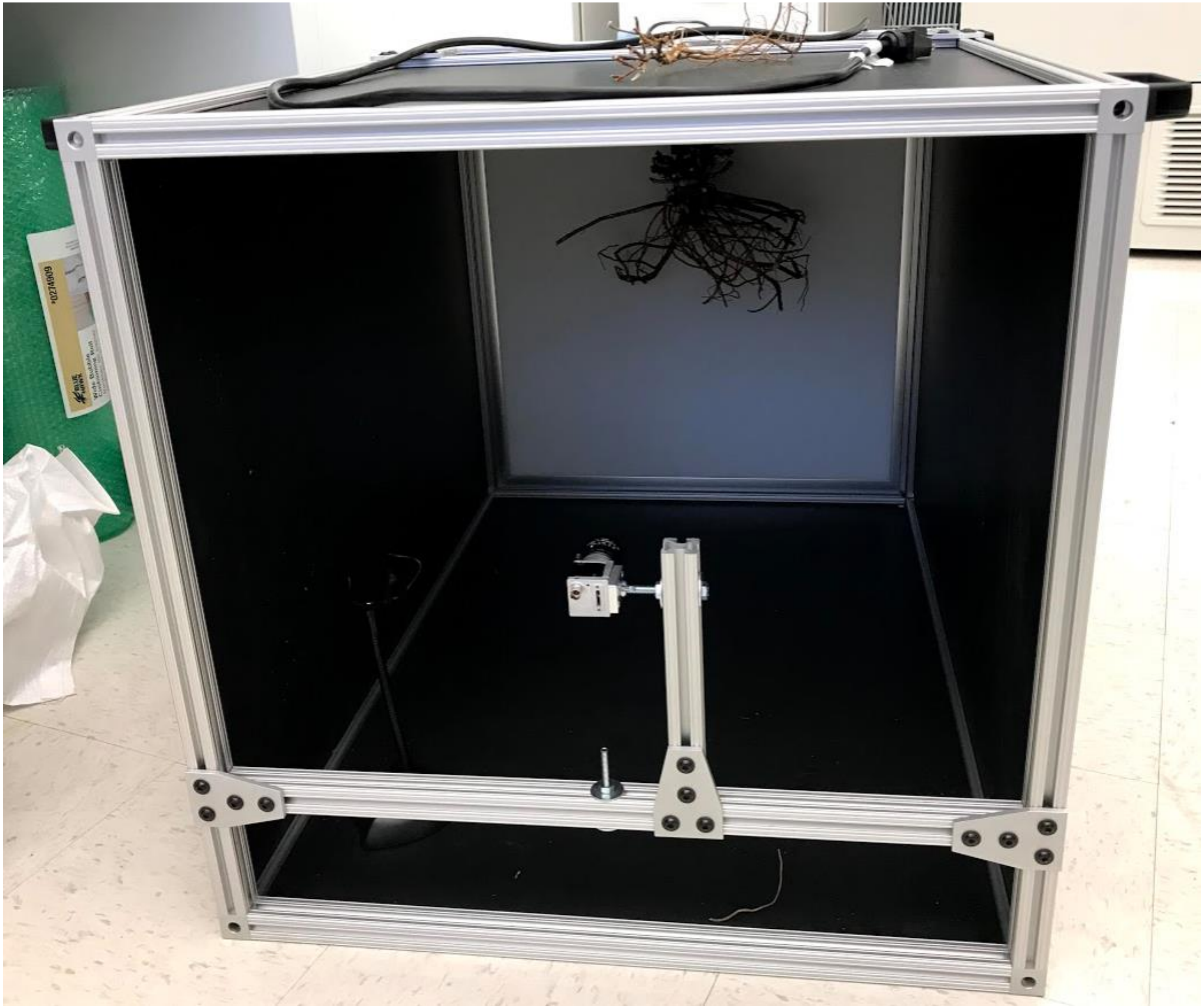

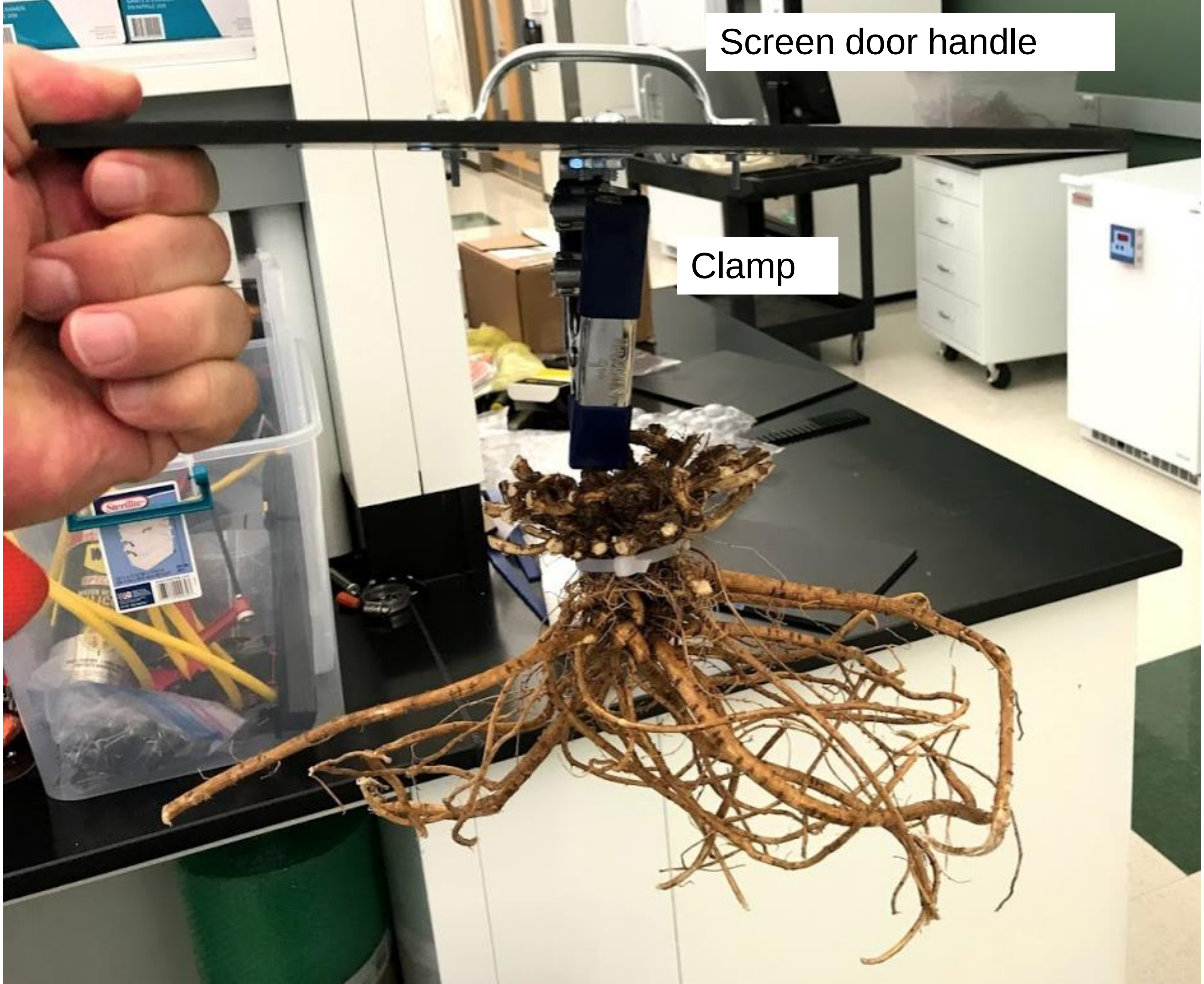
Screen door handle
Clamp

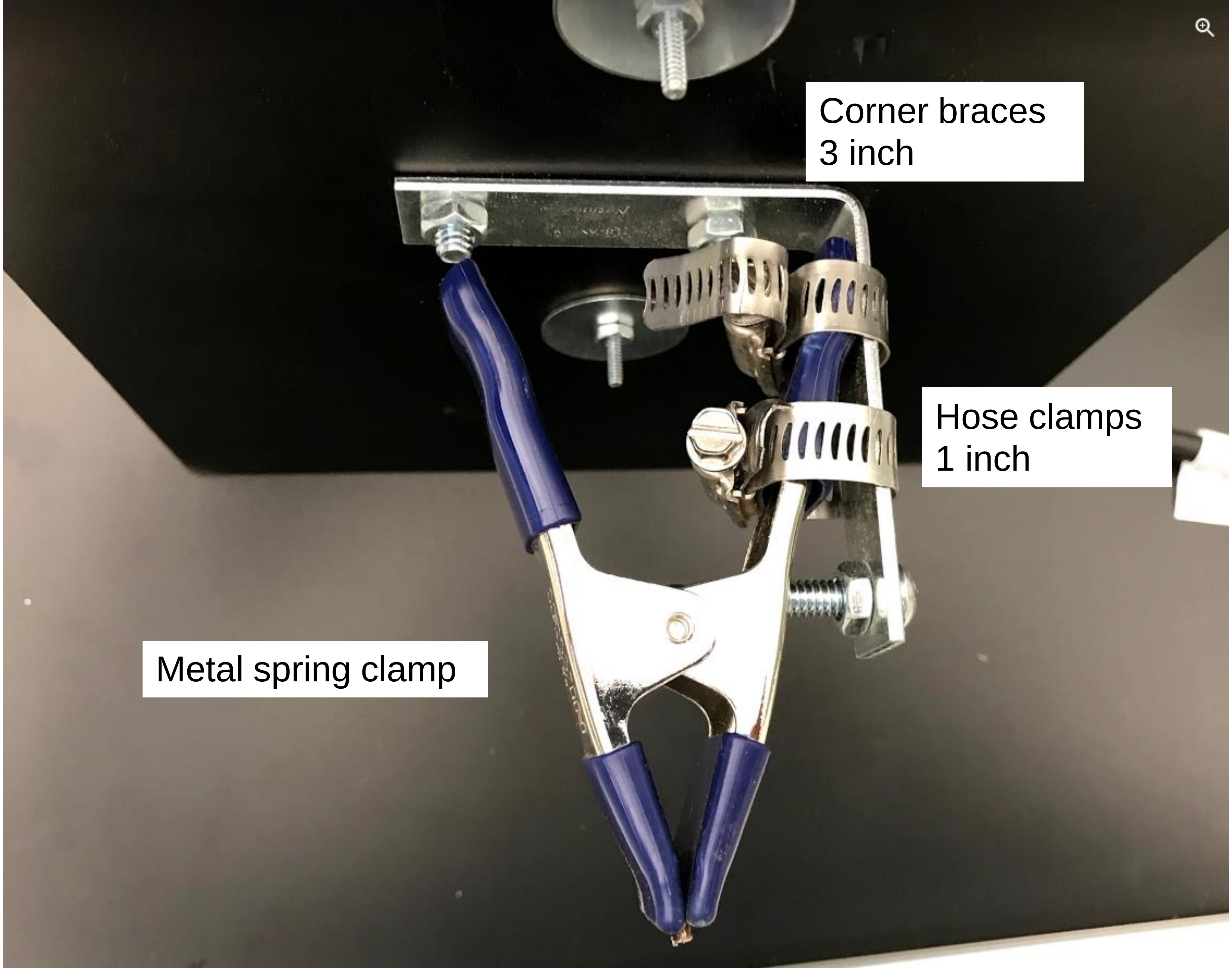
Corner braces
3 inch
Hose clamps
1 inch
Metal spring clamp

These ‘guide’ panels are glued to the side so the roots are always positioned correctly, dimensions give in PDF

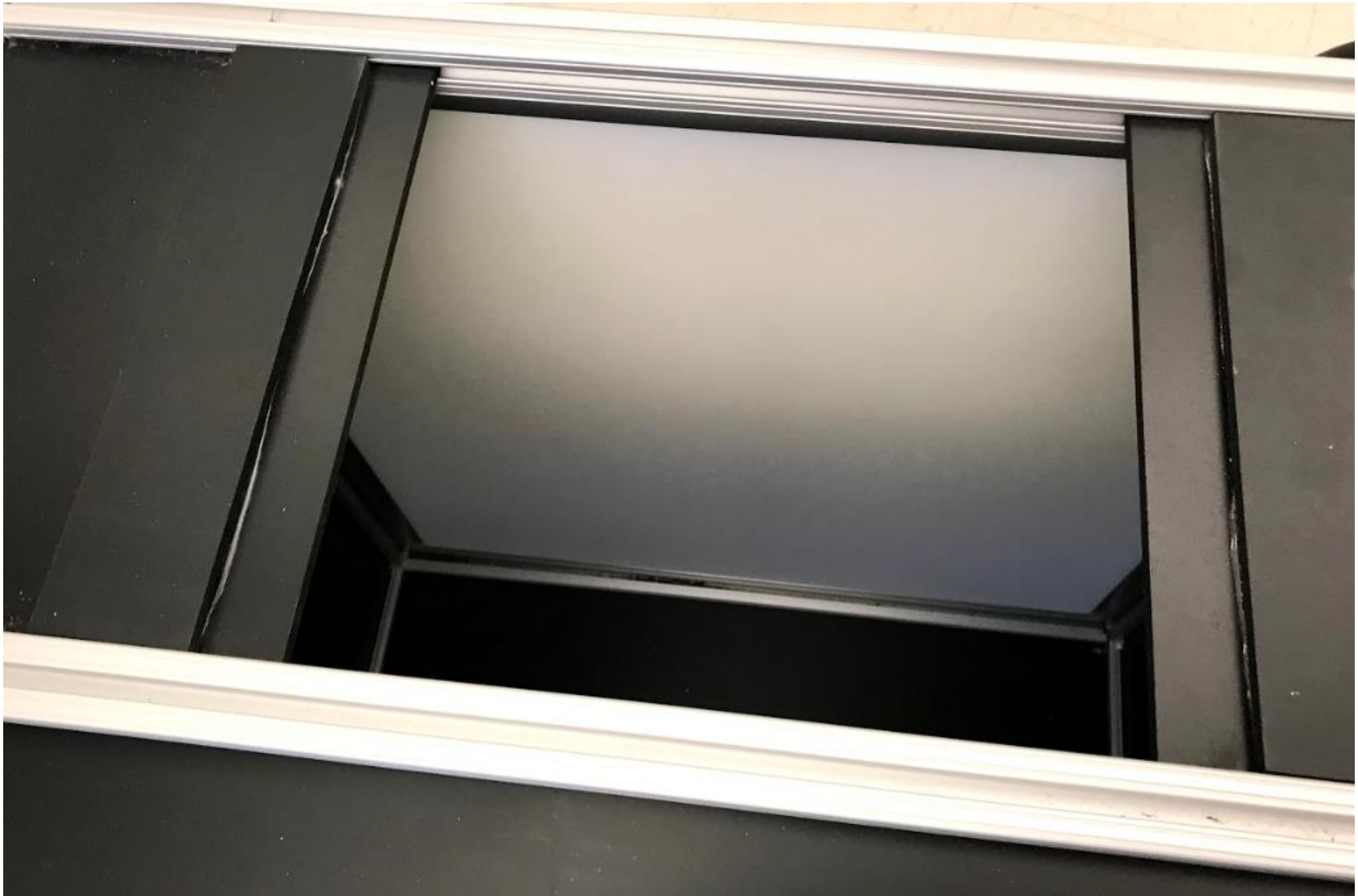

### RhizoVision Crown

Revision 4, adjust plastic panel sizes and cross profiles

Larry York, Noble Research Institute

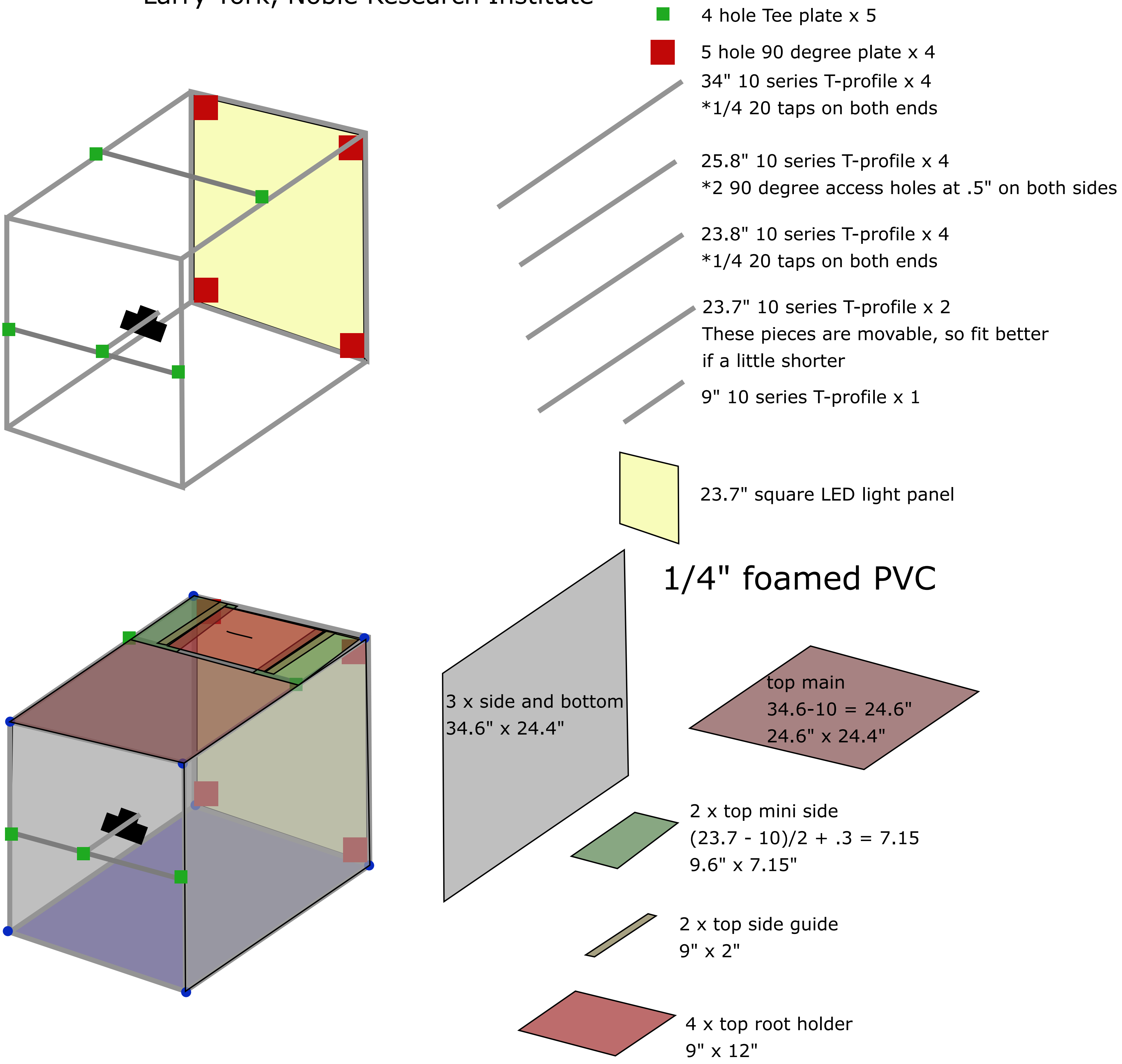

### Shopping Cart

Proceed to Checkout →

Continue Shopping

**Please visit the Shipping Schedule for more information**

| Image | Product Name | Machining Services / Each | Price | Qty | Subtotal | Remove |
|---|---|---|---|---|---|---|
| 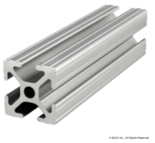<br>**SKU: 1010**<br>Move to wishlist | **34 Inches (863.6 Millimeters) 1.00" X 1.00" T-Slotted Profile - Four Open T-Slots**<br><br>**Part Number: 1010** | **1 Cut to Length**<br>**2 End Taps**<br>7061 Left<br>7061 Right | **$13.67** | 4 update | **$54.68** | Remove Item |
| 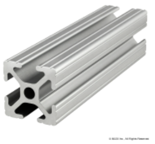<br>**SKU: 1010**<br>Move to wishlist | **25.9 Inches (657.86 Millimeters) 1.00" X 1.00" T-Slotted Profile - Four Open T-Slots**<br><br>**Part Number: 1010** | **1 Cut to Length**<br>**4 Access Holes**<br>7051 in R @ 0.5<br>7051 in R @ 25.4<br>7051 in S @ 0.5<br>7051 in S @ 25.4 | **$15.71** | 4 update | **$62.83** | Remove Item |
| 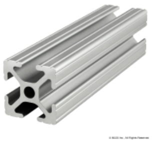<br>**SKU: 1010**<br>Move to wishlist | **23.9 Inches (607.06 Millimeters) 1.00" X 1.00" T-Slotted Profile - Four Open T-Slots**<br><br>**Part Number: 1010** | **1 Cut to Length**<br>**2 End Taps**<br>7061 Left<br>7061 Right | **$11.35** | 4 update | **$45.39** | Remove Item |
| 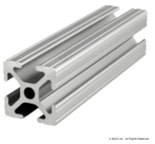<br>**SKU: 1010**<br>Move to wishlist | **23.9 Inches (607.06 Millimeters) 1.00" X 1.00" T-Slotted Profile - Four Open T-Slots**<br><br>**Part Number: 1010** | **1 Cut to Length** | **$7.45** | 2 update | **$14.89** | Remove Item |
| 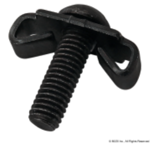<br>**SKU: 3381**<br>Move to wishlist | **Standard End Fastener, 1/4-20**<br><br>**Part Number: 3381** | None | **$1.50** | 16 update | **$24.00** | Remove Item |
| 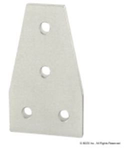<br>**SKU: 4141**<br>Move to wishlist | **10 Series 4 Hole - Tee Flat Plate**<br><br>**Part Number: 4141** | None | **$5.80** | 4 update | **$23.20** | Remove Item |
| 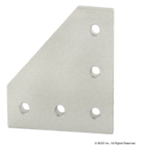<br>**SKU: 4151**<br>Move to wishlist | **10 Series 5 Hole - 90 Degree Angled Flat Plate**<br><br>**Part Number: 4151** | None | **$6.30** | 4 update | **$25.20** | Remove Item |
| 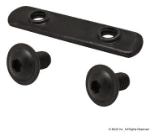<br>**SKU: 3356**<br>Move to wishlist | **Bolt Assembly: (2) 1/4-20 x .500" Black FBHSCS and Double Slide-In Economy T-Nut - Black Zinc**<br><br>**Part Number: 3356** | None | **$0.95** | 16 update | **$15.20** | Remove Item |

Update Shopping Cart

| | |
|---|---|
| Subtotal: | $265.39 |
| **Grand Total** | **$265.39** |

Code Apply Coupon

Proceed to Checkout →

**Please visit the Shipping Schedule for more information**
